# The Crunchometer: A Low-Cost, Open-Source Acoustic Analysis of Feeding Microstructure

**DOI:** 10.1101/2025.07.25.666891

**Authors:** Elvi Gil-Lievana, Benjamin Arroyo, Jesús Pérez-Ortega, Axl Lopez, Luis Alfredo Rodriguez Blanco, Xarenny Diaz, Gustavo Hernandez, Alam Coss, Emily Alway, Naama Reicher, Enrique Hernández Lemus, Maya Kaelberer, Diego V. Bohórquez, Ranier Gutierrez

## Abstract

Elucidating the neuronal circuits that govern appetite requires precise, high-resolution monitoring of the microstructure of solid food consumption, a need unmet by existing tools, which are either costly or lack the temporal resolution to align feeding events with neuronal activity. To overcome this, we developed the Crunchometer, a low-cost, open-source acoustic system that uses computational algorithms to generate high-resolution feeding ethograms from the sounds produced during solid food consumption. Validation across energy states (hunger/satiety) confirmed its sensitivity to changes in feeding microstructure, and the system reliably detected semaglutide-induced suppression of intake and reduced preference for a high-fat diet. Leveraging its seamless integration with in vivo recordings in freely behaving mice, we paired the Crunchometer with lateral hypothalamus (LH) electrophysiology to identify “meal-related” neurons that track entire meals rather than individual bouts. Calcium imaging further revealed that distinct subsets of LH GABAergic and glutamatergic neurons were tuned to feeding only, to licking only, or to both behaviors. Thus, LH neuronal ensembles differentially encode the consumption of solid food versus liquid sucrose. These findings demonstrate that the Crunchometer is a robust, accessible platform for dissecting the neural correlates of feeding behavior at the resolution of a single bite.

**Significance statement:** The Crunchometer is a low-cost, open-source technology that democratizes feeding analysis, enabling the precise dissection of neuronal circuits involved in appetite to advance therapies for obesity and eating disorders.

## Introduction

For decades, neuroscience research on appetite, taste, and reward has relied heavily on liquid food models, often reducing the study of ingestive behavior to simple measures of licking (Davis, 1989; Davis and Smith, 1992). The reasons for this are largely practical: liquids are easy to deliver with precision in experimental settings, such as fMRI scanners in humans (Smeets et al., 2019), and are well-suited for basic research on rodent studies (Gutierrez et al., 2006; Tellez et al., 2012). This methodological convenience, however, has fostered a narrow view of eating, oversimplifying the complex sensory and motor challenges unique to solid food. In rodents, eating solid food is a fundamentally different neurobiological event than sipping a liquid. The process begins with motor actions, such as biting and mastication (chewing), which in turn trigger a complex cascade of trigeminal (touch and texture) and gustatory (taste) stimulation. This is not a fixed mechanical action, but rather a dynamic, fine-grained sensory-motor feedback loop (Jacquin and Zeigler, 1983). Within this loop, sensory properties such as taste, texture, temperature, and hardness generate a rich stream of information known as “mouthfeel,” which is continuously relayed to the brain to influence eating rate, bite size, and ultimately, satiety (Gutierrez and Simon, 2021). Despite these critical differences, neuroscience has scarcely explored the neuronal correlates of the microstructure of solid food, focusing on liquid diets, operating under the largely untested assumption that both food forms recruit the same neuronal circuits, see (Pilato et al., 2024; Yamamoto et al., 1989), for exceptions. This lack of knowledge is mainly due to the high cost of current technologies and limitations of traditional techniques for monitoring food intake in laboratory animals.

Current methods to monitor feeding behavior could be classified into five different classes: 1) *Manual Weighing*: The most traditional and straightforward method involves manually weighing the food hopper or a dish of food at regular intervals. 2) *Automated Weighing Systems and Home Cage Monitoring Systems (Smart Cages)*: These systems build upon the engineering prowess of Curt Richter. In 1922, by continuously monitoring the food intake and activity of rats, Richter discovered the homeostatic functions of ingestive behavior (Richter, 1922). This equipment provides continuous monitoring using automated weighing scales, such as load cells, or other methods, such as the Snacker Tracker (Mueller et al., 2025). Home cage monitoring systems are the current state-of-the-art for tracking feeding behavior in a minimally invasive and ethologically relevant environment. Commercial systems, such as the BioDAQ (Research Diets) (Farley et al., 2003; Yang et al., 2025), OxyletProTM System (PANLAB, Cornellà, Spain) (Mariné-Casadó et al., 2018), Oxymax CLAMS-HC (Columbus Instruments), Promethion metabolic cage system (Sable Systems International, Las Vegas, NV, USA) (Ye et al., 2024), TSE PhenoMaster/IntelliCage system (TSE Systems, Bad Homburg, Germany) (Bake et al., 2014), PhenoTyper (Noldus IT) (Acosta-Rodríguez et al., 2017; Robinson and Riedel, 2014), employ sophisticated (and costly) automated weighing scales. These systems achieve high temporal resolution, enabling the analysis of meal patterns defined by bouts of grams consumed (Farley et al., 2003). The IntelliCage system excels in simultaneously monitoring the feeding behavior of multiple mice. However, it is not compatible with electrophysiological or calcium imaging recordings. Food intake in social contexts is a more ethologically valid model, in which radio-frequency identification (RFID) transponders enable the simultaneous assessment of feeding behavior across multiple mice co-housed in a single cage (Rathod and Fulvio, 2021). Still, they often find it challenging to precisely distinguish actual consumption from spillage or other feeding-related behaviors, such as biting, gnawing, licking, and locomotion. 3) *Automated Pellet Dispensers*: Often integrated into operant conditioning chambers, these devices provide a controlled way of delivering food pellets. While devices like the open-source Feeding Experimentation Device (FED3) (Ali and Kravitz, 2018; Matikainen-Ankney et al., 2021), a pellet dispenser, are useful for measuring reinforcement, they alter the natural feeding patterns of mice. For example, requiring a simple action, such as a nose-poke, can reduce overeating and weight gain in mice (Barrett et al., 2025). A further limitation is that FED3 may overestimate consumption if an animal retrieves and registers a pellet without actually consuming it. A significant strength of this method is its ability to enable closed-loop optogenetic stimulation concurrent with neuronal recordings. 4) *Video-Based Analysis*: While often used in conjunction with other methods, video analysis can provide unique insights into feeding behavior, but requires large video storage capacity and computer power, and frequently fails to distinguish crossing from a food zone from actually eating (Jennings et al., 2015). Moreover, direct human observation is labor-intensive and inherently subjective, compromising data quality.

5) *Audio-based methods* have been largely unexplored and underutilized (Ali and Kravitz, 2018), as they have mainly focused on using expensive microphones to detect rodents’ ultrasonic vocalizations (USVs) (Champeil-Potokar et al., 2023; Wardak et al., 2024), such as the 40 kHz vocalization associated with rat food consumption (Champeil-Potokar et al., 2023) or the 50 kHz vocalization linked to positive reinforcement (Wardak et al., 2024). Their inconsistent occurrence limits the utility of USVs, as they are not observed with every bite or in all subjects. Beyond these technical limitations, behavioral complexities further hinder accurate quantification. A further complication arises from the common hoarding behavior observed in mice. This practice of transporting food to other locations poses a considerable challenge to the precise assessment of spillage and the quantification of consumption. Consequently, most research has relied on liquid diets (e.g., sucrose or Ensure), in which lickometers enable precise, easy measurement of licking microstructure (Gutierrez et al., 2006; Spector et al., 1998; Tellez et al., 2012; Zhu et al., 2025). In contrast, the microstructure of eating solid food and its neuronal correlates remain poorly understood (Décarie-Spain et al., 2025), primarily due to technical limitations that impede detailed analysis of bite count and food chewing in mice (Stuber et al., 2025). To overcome these challenges, we developed the “Crunchometer,” a novel, cost-effective acoustic system for monitoring feeding. Unlike methods that rely on expensive microphones for USV detection, the Crunchometer utilizes an economical condenser microphone (Fifine K669 Amplitank, priced under $63) to analyze the temporal dynamics of feeding behavior by extracting and identifying bite sounds. This will enable scientists to create detailed feeding ethograms, thereby establishing an acoustic-based method to investigate the microstructure of solid food intake. This study introduces and validates the Crunchometer, a novel, cost-effective acoustic system that precisely analyzes the microstructure of solid food intake in mice. We demonstrate its utility by comparing automated bite detection with human observation, monitoring feeding patterns in various physiological states (hunger/satiety), characterizing pharmacological effects (semaglutide), and differentiating distinct feeding-related behaviors such as gnawing (i.e., biting an inedible object) vs. consumption induced via chemogenetics (Roth, 2016). Furthermore, we highlight its seamless integration with *in vivo* multichannel electrophysiology and microendoscope calcium imaging to elucidate the neural correlates of solid vs liquid food consumption.

## Results

### The Crunchometer: An Open-Source Sound-Based Method for Studying the Microstructure of Feeding Behavior

The setup of the Crunchometer is shown in **Figure 1A**, and an exploded view of its components is presented in **Figure 1B**. The Crunchometer consists of an acrylic box (1) with 11 holes drilled on one lateral wall (see Inset **Figure 1A**). During each session, two food pellets (one HFD (2) and one standard Chow (3)) were placed on a wall near the condenser microphone (4). A bottom-view camera (or top-view) continuously records the mice (5). Additionally, a 10% sucrose solution was dispensed to ensure mice had *ad libitum* access to both food and liquid. A Med Associates contact lickometer (6) could measure each lick and dispense a 1 μL drop of sucrose per lick (7). Videos were captured via Open Broadcaster Software (OBS) (OBS Project, 2025). The .mkv format was preferred over .mp4 due to its greater robustness and reliability during extended recording sessions, particularly in preventing data loss in case of power interruptions.

**Figure 1.**
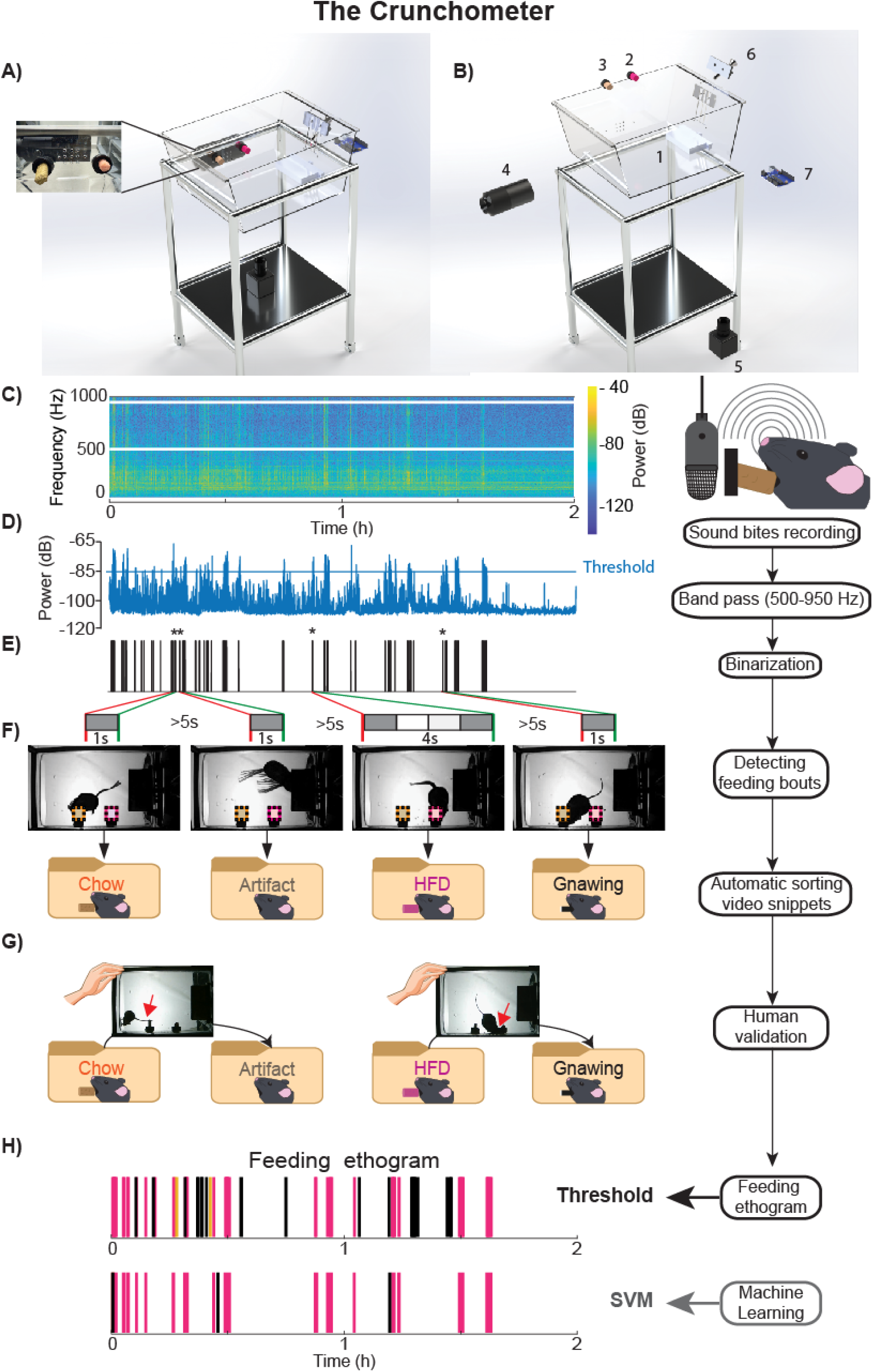
The Crunchometer System and Workflow for Acoustic Feeding Analysis. **A)** Schematic of the Crunchometer setup for behavioral feeding studies. This diagram illustrates the system’s integrated components for the acquisition and analysis of acoustic feeding data in mice. The inset details the microphone’s optimized placement, within the outer wall of the 11-hole box, and the precise locations of the food pellets. **B)** Exploded view of Crunchometer components. The acrylic box (1) in which a High-fat diet (HFD) (2) and Chow (3) pellets were positioned on the inner wall of the behavioral box (See **Video 1** for pellet preparation), at the same height as the condenser microphone (4). The microphone is mounted on the adjacent outer wall. A bottom-view camera (5) is installed beneath the box to record mouse locomotor activity (an optional top-view camera is not shown). A contact lickometer (6), controlled by an Arduino device or MedAssociate system (7), provides 1 µL of 10% sucrose per lick. **C)** Sound recording and processing workflow. Audio was sampled at 44.1 kHz, and video was recorded at 30 frames per second (fps). The spectrogram (1 s resolution) is also shown, with a color bar indicating power in dB. **D)** Power spectrum filtering. The power spectrum of the recorded audio was averaged over 500-950 Hz band to isolate potential bite sounds. The average dB in this band is plotted. A fixed threshold of -85 dB (determined by trial and error) was used in our setup. Every 1 s bin exceeding the threshold was assigned a value of 1; otherwise, it was assigned a value of 0. **E)** Identification of bite frames and feeding bouts. Black lines indicate putative bites, obtained through signal binarization and labeled as bite frames. Asterisks mark representative examples of feeding bouts, with red lines indicating bout onset and green lines indicating bout offset. Gray boxes represent 1-second time bins. Each feeding bout consists of one or more bites, separated by pauses of less than 5 seconds. Feeding bouts were detected within two defined Regions of Interest (ROIs) on the video recording, where the area corresponding to each pellet was outlined (yellow box for Chow, pink box for HFD). **F)** After identifying feeding bouts, the software automatically classified each video snippet by sorting them into four folders: “Chow,” “HFD,” “Gnawing,” or “Artifact.” The primary classification was determined by motion energy within two regions of interest (ROIs) drawn over the food: a yellow square for the Chow pellet and a pink square for the HFD pellet. For example, if motion energy was greater in the Chow ROI, the snippet was classified as ’Chow.’ Snippets were labeled ’Gnawing’ if visual inspection showed mouth movements without food consumption, such as biting the plastic cap that holds the pellet (or any other non-edible object). Finally, a snippet was labeled as ‘Artifact’ if a noise occurred while the mouse was outside both food ROIs. **G)** Human validation is a critical step to correcting automated classification errors. For example, the system may misclassify a snippet as “feeding” simply because the mouse is positioned within a feeding region of interest (ROI), even if it isn’t eating. The examples highlighted by the red arrows show such cases. During the manual review, a user corrects these errors by moving the misclassified snippets from the diet folders (Chow or HFD) to their proper category, such as “artifact” or “gnawing.” **H)** The final ethogram displays feeding bouts identified by two methods: the supervised Threshold method and a Support Vector Machine (SVM). In the graph, colored lines represent feeding on Chow pellets (yellow), HFD pellets (pink), and gnawing behavior (black). While the SVM method is more efficient at distinguishing bite-like sounds and less prone to artifacts, this precision comes at the cost of underestimating gnawing behavior. In **Supplementary** Fig. 1**-1**, the bill of materials is provided. **Supplementary** Fig. 1**-2** shows the robustness of Crunchometer bite detection to additive white noise. The Threshold method is sensitive to loud sounds overlapping the bite-related frequency band (500–950 Hz) but remains more reliable than the SVM method across the tested signal-to-noise ratio (SNR) range.

Before the feeding sessions, mice were habituated to the behavioral box over two consecutive days with 30-minute sessions to reduce stress. During the feeding session, synchronized video and audio streams were recorded in a single .mkv video file. The audio was subsequently extracted and stored in .ogg format for efficient processing, using the Crunchometer software (Tab 1(Processing Tools). 1. Audio Extractor). Potential sound bite events were identified by detecting the amplitude of sound frequencies between 500 and 950 Hz (**Figure 1C**). The power spectrum was averaged within this frequency band to isolate a potential sound bite. The audio snippets with frequencies within this interval were binarized: audio frames identified as putative bites were labeled “1,” while non-bite frames were labeled “0.” To improve bite detection accuracy, we calculated the mean power in the frequency band and applied a fixed threshold of -85 dB (**Figure 1D**). Although the optimal dB threshold can vary across setups, we utilized a fixed threshold calibrated for each setup. This approach was chosen because, unlike dynamic thresholds (e.g., z-score and 3 standard deviations above average noise), a fixed threshold more accurately detects when a mouse is not biting at all, such as during a state of satiety (note that Crunchometer software also lets you set a custom threshold or use a dynamic one using std; Tab 2 (Audio Detection)). Using these binary-labeled sound bites, we extracted the corresponding video frames (termed “video snippets”) for detecting feeding bouts. Given that microphones more reliably detect the abrupt noise of a broken pellet (i.e., a bite) but do not consistently capture chewing sounds, we opted to define feeding bouts to incorporate chewing times whenever possible. Accordingly, feeding bouts were defined as sequences containing one or more bites with Inter-Bout-Intervals (IBIs) of less than 5 seconds. Pauses longer than 5 seconds marked the start of a new feeding bout (**Figure 1E**). Using the onset and offset times of these video snippets, we automatically sorted them into the following behaviors: Artifact, Chow, HFD, and Gnawing (**Figure 1F**), based on the motion energy of two ROIs drawn over the pellets, which indicated whether the mice crossed either ROI. The Crunchometer, therefore, does not need to infer food identity acoustically: audio confirms that a bite occurred, and the mouse’s position within a food-specific ROI identifies which food was consumed. This design enables per-diet attribution even for pellets with indistinguishable crunch signatures. The final and most crucial step in our classification pipeline was human validation. A human observer supervised the automatic sorting of all video snippets. In the event of misclassification, snippets can be easily reclassified using the Crunchometer visual validator (Tab 3. (Snippets_ROIs) on the Crunchometer software). Common misclassifications occurred when the mouse’s tail touched a pellet or, more frequently, when it gnawed on a non-food object such as the plastic lid. This human validation was essential for ensuring the high fidelity of our behavioral database and mitigating the inherent limitations of automated classification. Finally, an ethogram was constructed to visualize the temporal distribution of these behaviors throughout the session. This was achieved using both supervised thresholding and an unsupervised Support Vector Machine (SVM) approach (**Figure 1G**). The Crunchometer successfully captured individual sound bites from mice, enabling us to construct detailed ethograms of feeding behavior on Chow and HFD pellets and gnawing behavior (**Figure 1H**). In summary, the Crunchometer enables the systematic detection of individual bites, providing a robust platform for monitoring complex feeding dynamics.

### The Crunchometer was more precise and reliable than humans in detecting the start and end of a feeding bout

After defining the Crunchometer’s signal processing, we evaluated our sound-based feeding bout detection method by comparing its performance with that of human observers to identify key differences. We analyzed the feeding behavior of six mice using the Crunchometer. Separately, seven human observers independently annotated feeding bouts using the open-source software BORIS (Behavioral Observation Research Interactive Software) (Friard and Gamba, 2016). Both the Crunchometer and human observers consistently detected feeding bouts (**Figure 2A**, top panel). However, some feeding bouts were more challenging to detect and were identified only by the Crunchometer or human observers (see black arrows in the bottom panel of **Figure 2A**). We quantified the agreement between human observers and the Crunchometer using Normalized Mutual Information (NMI). This metric, which ranges from 0 (no overlap) to 1 (100% similarity), measures their shared information. Initially, four of the seven human observers (Humans 2, 3, 4, 6; pairs human 3 vs. 6 (NMI = 0.53), 3 vs. 4 (NMI = 0.55), 2 vs. 4 (NMI = 0.61), and 2 vs. 3 (NMI = 0.64)) achieved higher agreement NMI values than the remaining three humans (7, 5, and 1). This variability within humans likely reflects limitations in attention span, reaction times, or fatigue, potentially leading to delayed or missed detection of feeding bouts (**Figure 2B**). Overall, the within-human agreement achieved a mean ± standard deviation of NMI = 0.38 ± 0.14. In contrast, both the Threshold and SVM methods exhibited greater similarity in their ability to detect feeding bouts (**Figure 2B**; NMI = 0.58). However, the similarity between human and Crunchometer was consistently lower compared to within-group comparisons (**Figure 2B**, blue rectangle; max = 0.36, mean ± std = 0.27 ± 0.05), highlighting notable differences in how the Crunchometer and human observers identified feeding bouts.

**Figure 2.**
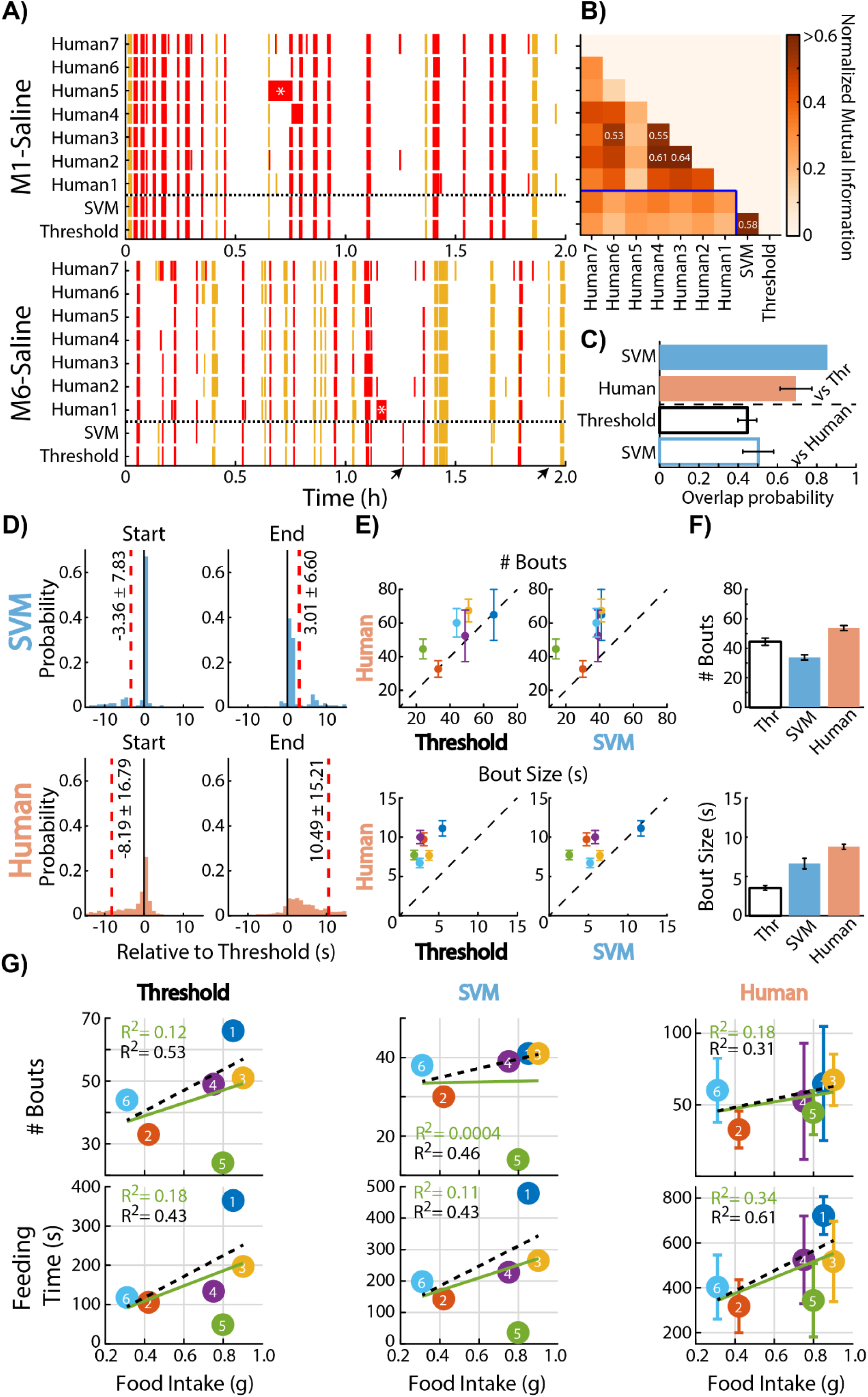
The Threshold and SVM methods were more precise and reliable than humans at detecting the start and end of a feeding bout; however, these methods tend to fragment feeding bouts more than humans do, as they estimate more events and longer feeding periods. **A)** Representative ethograms illustrate the feeding behavior of two fasted mice. Red and yellow marks denote feeding bouts related to HFD and Chow pellets, respectively. A horizontal dashed black line separates human-annotated feeding bouts from method-detected feeding bouts. The experiment in the top panel (M1-Saline) demonstrates greater consistency between human observations and Crunchometer detections, whereas the black arrows in the bottom panel (M6-Saline) highlight two feeding bouts that were uniquely detected by either mathematical methods or by human observers. Also, the asterisk (*) indicates periods in which a human failed to detect a feeding bout correctly. **B)** To evaluate the similarity between the automated methods and human detections, Normalized Mutual Information (NMI) was used, with varying shades of orange indicating different NMI values. While both human observers and the methods showed high within-group correlation, there was notably reduced similarity between human and method detections (highlighted by the blue rectangle). **C)** To assess the overlap probability between the Crunchometer methods and human observers, we first concatenated all feeding bouts detected across the six experiments. Subsequently, we quantified the total number of feeding bouts identified by human observers and by each Crunchometer method. Solid bars represent the overlap probability that an SVM and seven human observers agreed on the detection of a single feeding bout, relative to the 267 total bouts detected by the Threshold method across all six mice. Empty bars display the overlap probability between Threshold and SVM detections relative to each of the seven human observers (with total feeding bouts detected by each observer being 251, 286, 228, 248, 583, 215, and 444 for human 1 through 7, respectively). Horizontal black lines indicate the mean ± SEM. **D)** Time delay in detections by SVM and Human observers relative to Threshold-based onsets. The probability distributions of detection times by the SVM (top panels) and Human observers (bottom panels) are shown relative to the start (left panels) and end (right panels) of Threshold-based detections. The solid black line indicates the Threshold onset, while the red dashed line depicts the mean detection delay. Delay values are presented as mean ± std. **E)** Scatter plots illustrate the number and size of feeding bouts detected by human observers and the Crunchometer methods. Points falling on the diagonal dashed line indicate similar detection rates between human and automated methods. Deviations below the line (lower right) suggest greater detection by the Crunchometer methods, while deviations above the line (upper left) signify more detections by human observers. The left panels compare human detections against the threshold-based method, and the right panels compare them against the SVM-based method. Errors indicate the mean ± SEM. **F)** Total number (top) and size (bottom) of feeding bouts detected by human observers and the Crunchometer methods. Bars represent the mean for each detection source, with error bars indicating the mean ± SEM. **G)** Pearson coefficient of determination, R-squared, of mouse intake with bout size and bout number from automated methods and human observers. These scatter plots display the correlation between food intake and either feeding size or the number of bouts across all experiments. The black dashed line represents the linear regression excluding one subject (mouse number 5), whereas the green solid line indicates the regression including all subjects. For human detections, error bars are the mean ± SEM.

To further explore the differences between humans and the Crunchometer, we examined the overlap in feeding bout detections across sessions. The SVM method demonstrated greater overlap with the Threshold method (an overlap probability above 0.8, indicating at least one bin overlaps) than with human observers (**Figure 2C**, top panel). This means that the SVM frequently identified the same moments as eating times as those identified by the Threshold method. Humans also agree with the feeding bout intervals detected by the Threshold method (overlap probability humans vs Threshold around 0.7). In contrast, both Crunchometer methods exhibited less overlap than human observers (**Figure 2C**, bottom panel, with overlap probabilities below 0.5 for both methods). This result indicates that the SVM and Threshold automated methods for detecting feeding bouts are more similar to each other compared to human observation.

We quantified the response times for detecting the bout onset and offset using both the SVM method and human observers, relative to the Threshold method (Time = 0 s). Human observers exhibited longer delays in identifying the feeding bouts (average response relative to Start: -8.19 s and End: +10.46 s) compared to the SVM method (Start: -3.36 s and End: +3.01 s; **Figure 2D**). Consequently, human observers exhibited lower precision in detecting both the onset and offset of feeding bouts than the more reliable performance of the Threshold and SVM methods.

Next, we quantified the number and size of bouts detected by Crunchometer methods compared to those detected by humans. The Threshold method detected a similar number of bouts as human observers, but these bouts had shorter durations (**Figure 2E**, left panels). In contrast, the SVM method identified fewer bouts with intermediate durations (**Figure 2E**, right panels). Overall, human observers detected more and longer feeding bouts than both Crunchometer methods. This was because the Threshold method tended to fragment feeding events, while the SVM method captured fewer bouts (**Figure 2F**).

We finally examined whether the number of bouts detected could predict the actual food intake (in grams) of the mice during the 2-hour feeding session. Initially, we observed a low Pearson coefficient of determination across all conditions between the number of feeding bouts and total intake (**Figure 2G**, top panel; see R² in green) or between bout size and total intake (**Figure 2G**, bottom panel; see R² in green). This lack of correlation stemmed from one subject (mouse 5), which consumed approximately 0.8 g but exhibited very few, short-duration bites, suggesting a highly efficient bite pattern. Notably, detecting feeding behavior in this specific mouse proved difficult for both the Crunchometer and human observers. Consequently, after excluding this mouse, we recalculated the correlations and found that the best predictor for food intake depended on the method used. When using the number of feeding bouts (**Figure 2G**, top panel), the Threshold method exhibited the strongest correlation with food intake (R² = 0.53, dashed black line), surpassing both the SVM method (R² = 0.46) and human observations (R² = 0.31). However, when using the total feeding time as the predictor (**Figure 2G**, bottom panel), human observers were most effective, achieving the highest correlation (R² = 0.61) compared to both the Threshold and SVM methods (R² = 0.43 for both). This highlights the fundamental distinction in detecting when a mouse eats: the Crunchometer excels at detecting discrete, sound-producing bites, making it a precise event counter. In contrast, by integrating both auditory and visual cues, such as quiet chewing and eating spilled crumbs, humans are better able to estimate total food intake over a continuous feeding period.

### The Crunchometer System Distinguishes Meal Patterns of Fed and Fasted Mice

We initially validated our novel food intake monitoring system, the “Crunchometer,” by measuring consumption in six naive wild-type mice under both fed and 18-hour fasted conditions. Behavioral observations were conducted over a two-hour timeframe, a duration sufficient to elicit the entire behavioral satiety sequence following feeding (Halford et al., 1998; Tejas-Juárez et al., 2014). A visual inspection of the feeding ethograms revealed that fasted mice exhibited a greater number of feeding bouts compared to their sated counterparts (**Figure 3A**). Notably, one mouse (number 5) displayed robust gnawing behavior, indicated by black ticks in **Figure 3A**, specifically while in a fasted state. Supporting these observations, the cumulative feeding time was significantly higher in the fasted group, nearly doubling that of the fed group (Two-sample Kolmogorov-Smirnov test: D = 0.92500, *p* < 0.0001; **Figure 3A**, bottom panel). As predicted by the behavioral satiety sequence (Halford et al., 1998), fasted mice initially exhibited short IBIs that progressively lengthened as they approached satiety. This pattern is illustrated in the upper panel of **Figure 3B**, which plots the average IBI in 10-minute increments. We observed a significant increase in IBI time bin at 1.16 h (Mann–Whitney U test: U = 2, *p* = 0.0317). Likewise, the feeding rate increased significantly within the first 10 minutes of the session (two-way ANOVA, time effect: F_(11,110)_ = 2.663, *p* = 0.0046, physiological state effect: F_(1,10)_ = 4.293, *p* = 0.0651, and interaction: F_(11,110)_ = 3.474, *p* = 0.0003, *post hoc* analysis: time bin = 0.17 h *p* <0.0001; time bin = 0.58 h *p* = 0.0349; **Figure 3B**, lower panel). Finally, we found a significant correlation between the total grams of food consumed and total feeding time for both fed and fasted mice (**Figure 3B**, right panel). This positive correlation indicates that longer feeding durations were associated with greater food intake.

**Figure 3.**
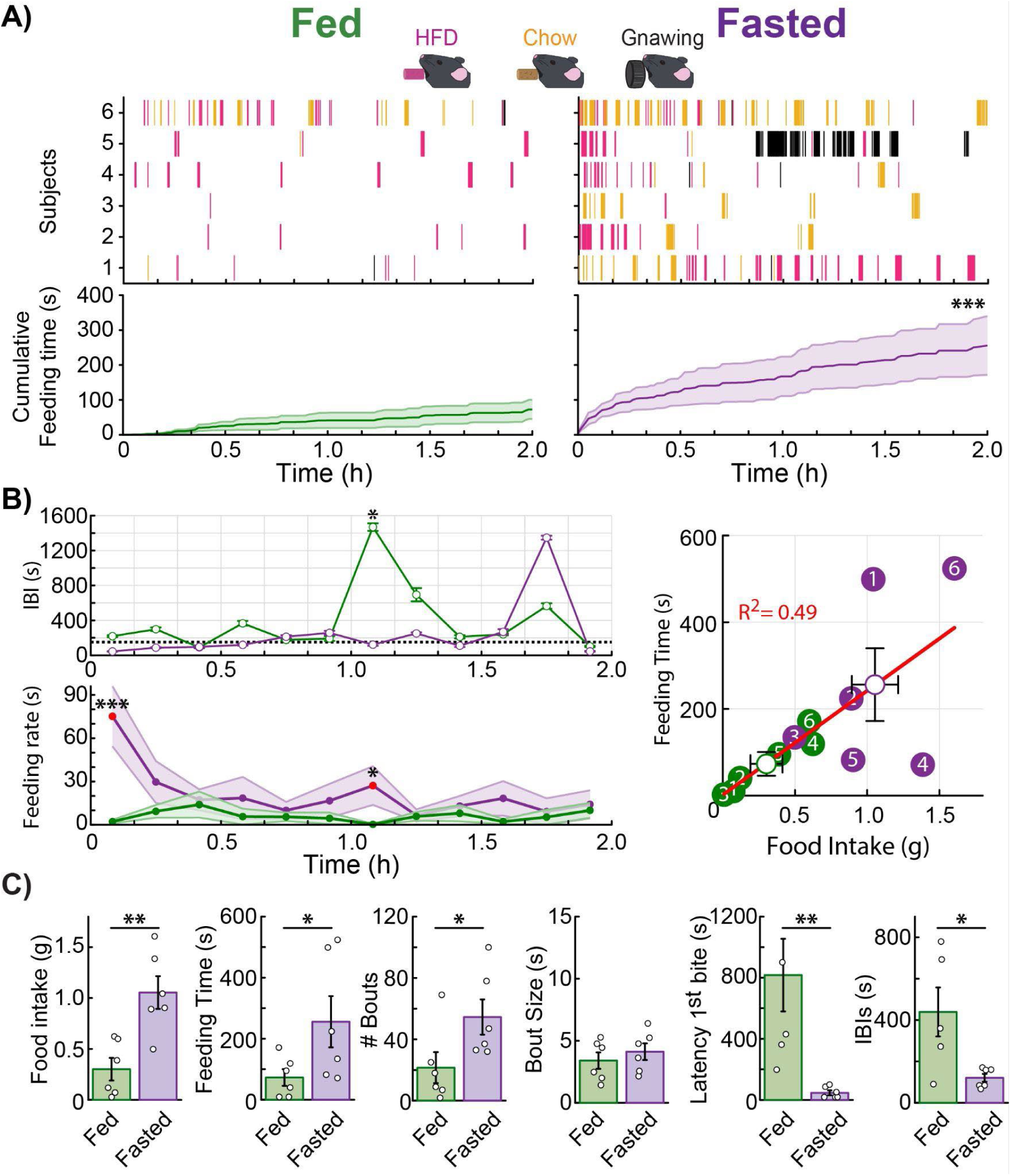
The Crunchometer captures detailed differences in feeding microstructure between satiated and fasted states. **A)** Representative feeding ethograms from a satiated (Fed) and a food-deprived (Fasted) mouse. Lines indicate bouts of HFD consumption (pink), standard Chow consumption (yellow), and non-edible gnawing (black). The lower panel shows the cumulative feeding time (only for Chow and HFD pellets, gnawing is excluded) over a two-hour session for the fed (green) and fasted (purple) groups (n = 6 mice). Shaded areas indicate the standard error of the mean (± SEM). **B)** The top panel shows the mean Inter-Bout Interval (IBI) in 10-minute bins, illustrating the progression toward satiety. The bottom panel shows the corresponding feeding rate; red dots indicate a significant difference between fed and fasted states. The scatter plot (right) shows a significant correlation between total feeding time and food intake (R² = 0.49, *p* = 0.011). **C)** Quantification of feeding parameters, including total feeding time, number of bouts, bout size, latency to first bite, and mean IBI. Data are expressed as mean ± SEM. \**p* < 0.05, \*\**p* < 0.01, \*\*\**p* < 0.001.

The Crunchometer could also quantify various behaviors related to the microstructure of feeding. These metrics include not only total food consumption (in grams) and feeding time (in seconds), but also the number and size of feeding bouts, latency to the first bite, and the IBI. As expected, total food intake was significantly different between the groups: fasted mice consumed more grams of food than fed mice (Mann–Whitney U test: U = 2, *p* = 0.0043). Likewise, compared to fed mice, fasted mice exhibited a significant increase in total feeding time (Mann–Whitney U test: U = 7, *p* = 0.0465; **Figure 3C**) and in the number of feeding bouts (Mann–Whitney U test: U = 4, *p* = 0.0130; **Figure 3C**). However, the average bout size did not differ significantly between the groups (Mann–Whitney U test: U = 12, *p* = 0.1970; **Figure 3C**). The latency to the first bite was significantly shorter in the fasted group, suggesting an increased motivation to eat (Mann–Whitney U test: U = 0, *p* = 0.0011; **Figure 3C**). Similarly, the IBIs across the entire session were significantly shorter in fasted mice than in fed mice (Mann–Whitney U test: U = 3, *p* = 0.0152; **Figure 3C**). Taken together, these results demonstrate that the Crunchometer can distinguish between the physiological energy states of hunger and satiety. These states were differentiated primarily by the overall feeding pattern (total feeding time), an increased initial feeding rate, and a higher number of bouts separated by short IBIs. In simple terms, hungry mice eat more frequently, especially at the beginning of the session.

### Semaglutide Suppresses Feeding and Reduces Preference for a High-Fat Diet

To characterize the acute anorexigenic effects of semaglutide, a widely used GLP-1 receptor agonist (Huang et al., 2024; Kim et al., 2024; Knudsen and Lau, 2019; Teixidor-Deulofeu et al., 2025), we used the Crunchometer to analyze feeding microstructure in detail. Mice were administered saline or semaglutide via subcutaneous injection immediately before the behavioral experiment began. Under baseline conditions (fasted, saline-treated), mice exhibited robust feeding with a clear preference for the HFD over standard Chow (Two-sample Kolmogorov-Smirnov test: D = 0.91667, *p* < 0.0001) (**Figure 4A**). In sharp contrast, following the administration of semaglutide (0.123 mg/kg) (Zhang et al., 2023), the same fasted mice displayed markedly suppressed feeding. Surprisingly, they also lost their strong preference for the HFD pellet (**Figure 4B**). This reduction in feeding behavior persisted in a follow-up test 24 hours later, even with *ad libitum* access to food (**Figure 4C**). We observed a significant, acute reduction in body weight at 24 h post-semaglutide administration (D1 Post-sem), with mice losing approximately 5% of their total mass (**Figure 4D**). This effect was restricted to the immediate post-administration period; from D2 to D5, the mice showed a steady recovery and continued their growth trajectory. By D5, body weight was significantly higher than the initial control levels (one-way ANOVA, F(6,30) = 11.55, *p* < 0.0001; post hoc analysis: Ctrl vs D1 Post-sem *p* = 0.0214, Ctrl vs D4 Post-sem p < 0.0099, Ctrl vs D5 Post-sem p < 0.0001; **Figure 4D**). Semaglutide treatment led to a significant decrease in total food intake and a sharp reduction in preference for HFD (two-way ANOVA, treatment effect: F_(2,30)_ = 26.88, *p* < 0.0001, diet effect: F_(1,30)_ = 16.74, *p =* 0.0003 and interaction: F_(2,30)_ = 23.53, *p* = < 0.0001; *post hoc* analysis: Ctrl Chow vs. Ctrl HFD, *p* < 0.0001; Sem Chow vs Sem HFD, *p* = 0.8731; Post sem Chow vs Post sem HFD p = 0.4857; **Figure 4E**). Likewise, mice spent significantly less feeding time under semaglutide treatment, and the next day, followed up test (two-way ANOVA, treatment effect: F_(2,30)_ = 16.96 *p* < 0.0001, diet effect: F_(1,30)_ = 0.7130, *p =* 0.4051 and interaction: F_(2,30)_ = 10.85, *p* = 0.0003; *post hoc* analysis: Ctrl vs. Sem, p = 0.0028; Ctrl vs Post sem *p* = 0.0019; **Figure 4E**, middle panel). Quantitative analysis of the feeding microstructure confirmed and extended these observations: Acute semaglutide administration significantly reduced both the number of feeding bouts (two-way ANOVA, treatment effect: F_(2,30)_ = 16.96, *p* < 0.0001, diet effect: F_(1,30)_ = 0.7130, *p =* 0.4051 and interaction: F_(2,30)_ = 10.85, *p* = 0.0003; *post hoc* analysis: Ctrl vs Sem *p* < 0.0001, Ctrl vs Post sem *p* < 0.0001; **Figure 4E**, right panel) and the size of each bout compared to saline-treated controls but only in the Post sem day (one-way ANOVA, F_(2,15)_ = 4.394, *p* = 0.0315; *post hoc* analysis: Ctrl vs Post sem *p* = 0.0097; **Figure 4F**). No significant difference was found in latency to the first bout. Consistent with the induction of satiety, the IBIs were significantly longer in mice treated with semaglutide (one-way ANOVA, F_(2,15)_ = 5.929, *p* = 0.0127; *post hoc* revealed a significant difference between Ctrl vs. Sem, *p* = 0.0036; **Figure 4F**). The anorexigenic effect of semaglutide extended to other palatable stimuli; the consumption of 10% sucrose solution was also significantly reduced (one-way ANOVA, F_(2,15)_ = 21.37, *p* < 0.0001; *post hoc* analysis: Ctrl vs Sem *p* < 0.0001, Ctrl vs Post sem *p* < 0.0001; **Figure 4F**). Finally, analysis across the session revealed that semaglutide-treated mice took significantly longer pauses between feeding bouts (IBIs) (Mann–Whitney U test: U = 0, time bin = 1.16 h Ctrl vs Sem, *p* = 0.0119; U = 0, time bin = 1.5 h Ctrl vs Post sem, *p* = 0.0357; U = 0, time bin = 1.66 h Ctrl vs Post sem, *p* = 0.0179; **Figure 4G**, left panel) and exhibited a correspondingly lower feeding rate compared to saline-treated controls (**Figure 4G**, middle panel). Additionally, feeding was significantly correlated with food intake (**Figure 4G**, right panel). These findings are consistent with the induction of a satiety-like state. Notably, the semaglutide-treated mice clustered with the satiated (fed) and post-semaglutide groups, indicating that they shared a similar low-feeding phenotype. In summary, these results demonstrate that acute semaglutide administration produces a powerful satiety-like state, characterized by fewer feeding bouts and longer IBIs. Furthermore, the Crunchometer analysis reveals that semaglutide also markedly reduces the preference for a palatable high-fat diet (**Video 2**). These findings validate the Crunchometer as a sensitive tool for dissecting the complex behavioral effects of anorexigenic drugs.

**Figure 4.**
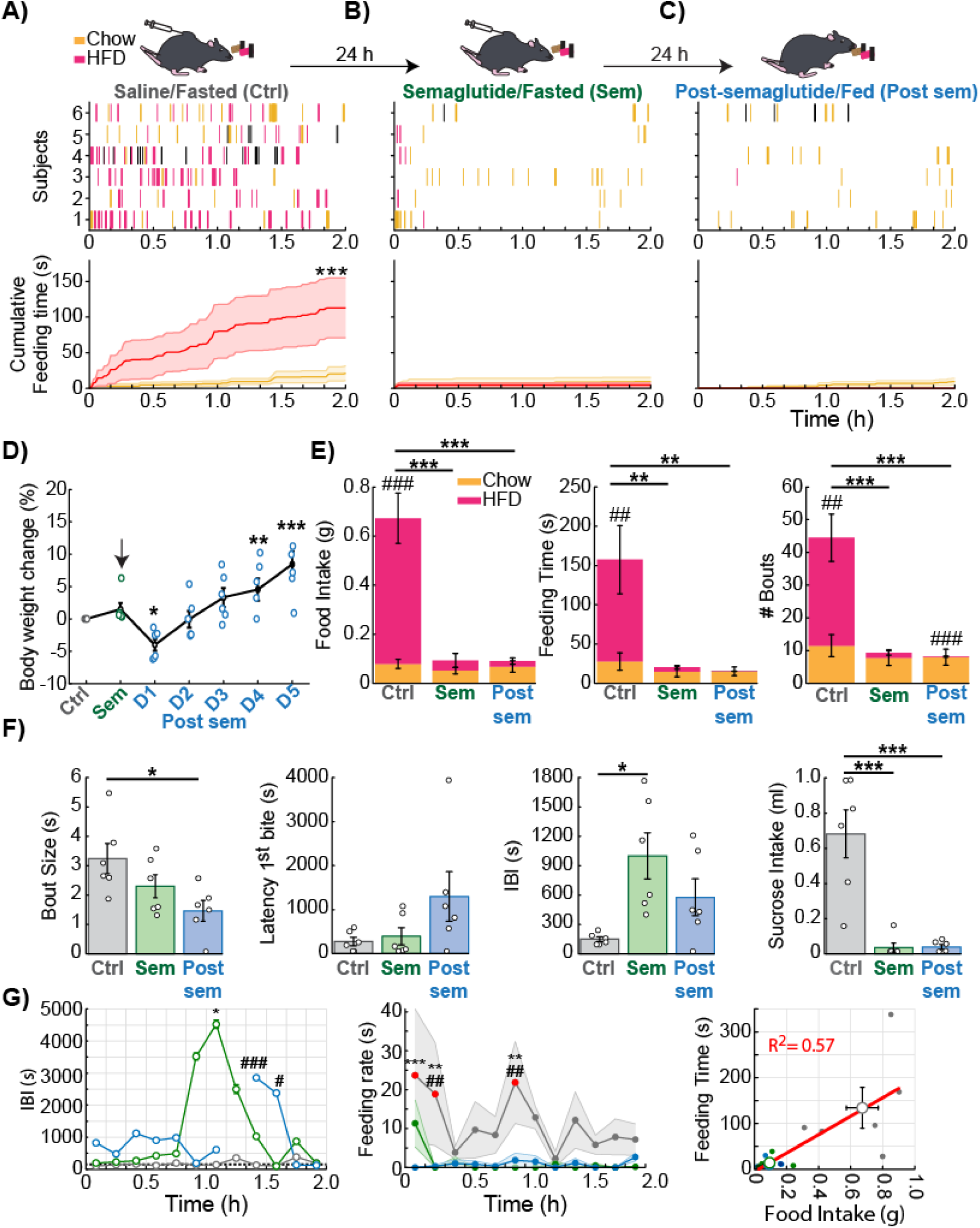
Semaglutide suppresses appetite and reduces the preference for a high-fat diet. **A)** Feeding bouts and cumulative intake in fasted mice following subcutaneous administration of saline (control group, Ctrl). **B)** The same parameters were measured in the same mice 24 hours later, under fasting conditions, following subcutaneous semaglutide administration (Sem group). **C)** Feeding behavior recorded 24 hours after semaglutide administration under fed conditions (Post-Sem group). In the ethograms, pink lines represent HFD pellet consumption, yellow lines indicate pellet consumption, and black lines denote gnawing events. In cumulative intake plots, solid lines show group averages (n = 6 mice), and shaded areas represent the ± SEM. All three experimental conditions (Ctrl, Sem, and Post-Sem) were tested sequentially in the same animals, with 24-hour intervals between sessions, as indicated by the arrow in the experimental timeline. **D)** Percentage change in the body weight of mice. The mice were weighed prior to the onset of the Crunchometer test. The black arrow indicates the administration of semaglutide. The percentage change was calculated by subtracting the body weight recorded on the saline administration day (Ctrl) from the weights on subsequent days (semaglutide and post-semaglutide), then dividing by the baseline (Ctrl) weight and multiplying by 100. Asterisks indicate statistically significant differences between Ctrl vs D1, D4 and D5 Post-sem. Data are expressed as mean ± SEM. *p < 0.05, **p < 0.01, ***p < 0.001. **E)** Bar plots of food intake, feeding time, and number of bouts for Chow pellet (yellow bars) and HFD pellet (pink bars) across the three experimental groups. The sum of the yellow and pink bars represents the total food intake. Asterisks indicate statistically significant differences between experimental groups (Ctrl (Sal/Fasted), Sem (Sem/Fasted), and Post sem (Post-semaglutide/Fed)), while the hash symbol (#) denotes significant differences between Chow and HFD pellets within the same group. **F)** Quantitative analysis of feeding variables measured using the Crunchometer: latency to the first bite, bout size, IBI, and intake of a 10% sucrose solution. Data are expressed as mean ± SEM. \**p* < 0.05, \*\**p* < 0.01, \*\*\**p* < 0.001. **G)** Average IBIs and feeding rate calculated in 10-minute bins across the session. For the feeding rate, red dots indicate a significant difference between groups (two-way ANOVA, time effect: F_(11,165)_ = 1.386, *p* = 0.1837, Treatment effect: F_(2,15)_ = 7.310, *p* = 0.0061, and interaction: F_(22,165)_ = 0.8404, *p* = 0.6721; *post hoc* analysis: Ctrl vs Post sem time bins = 0.17 h, 0.33 h, 1 h (*p* < 0.05); Ctrl vs Sem time bins = 0.33 h, 1 h (*p* < 0.05). Asterisks indicate statistically significant differences between Ctrl vs Sem (Sal/Fasted vs Sem/Fasted), while a hash symbol denotes significant differences between Ctrl vs Post sem (Sal/Fasted vs Post-semaglutide/Fed). The scatter plot in the right panel shows the correlation between feeding time and total food intake (R² = 0.57, *p* = 0.00031). A group analysis, as shown here, can be performed using the “short.mat” files generated by the Crunchometer software, via Tab 6 (Group analysis). The “short.mat” files need to be in a folder named after each experimental group.

### The Crunchometer Distinguishes Food Intake from Gnawing Behavior Induced by LH GABAergic Neuron Activation

We next sought to determine the effect of chemogenetic activation of GABAergic neurons in the LH on feeding behavior. To this end, an AAV8-hSyn-DIO-hM3D(Gq)-mCherry virus was bilaterally injected into the LH of Vgat-cre mice (n=4), allowing for the expression of the activating DREADD transgene over three weeks. The reporter protein mCherry was expressed in the neuronal somas of the LH, confirming localized transfection (**Figure 5A**). Under baseline satiety conditions, mice injected intraperitoneally with saline exhibited fewer feeding bouts than fasted mice; however, both groups showed a preference for the HFD pellet (**Figures 5B-C**). In contrast, chemogenetic activation of LH GABAergic neurons in satiated mice via CNO administration (Clozapine N-Oxide) led to a paradoxical behavioral pattern. It increased overall consummatory actions, particularly towards both Chow and HFD pellets, and exacerbated spillage and gnawing behavior (**Figure 5D**; see **Video 3**).

**Figure 5.**
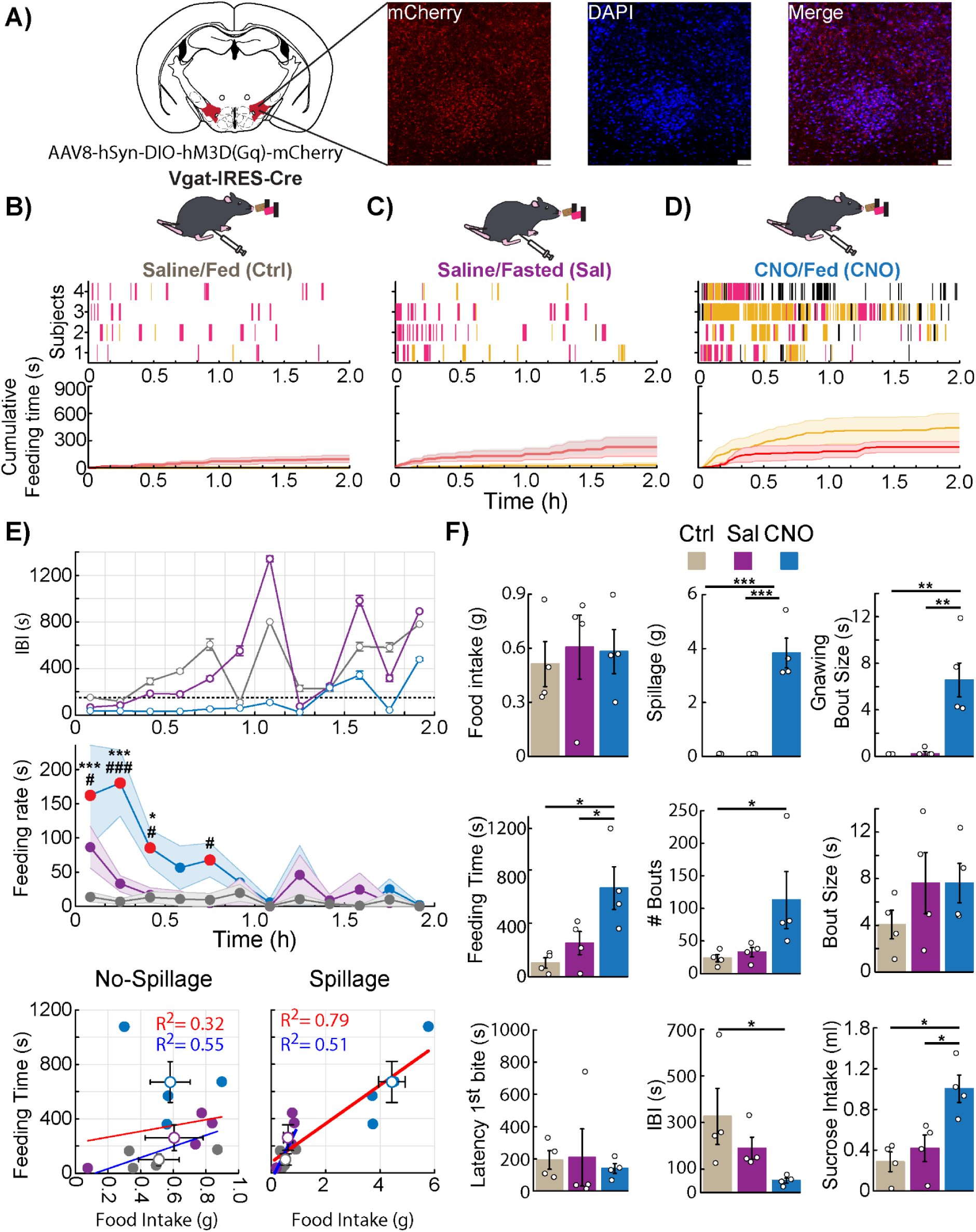
Bilateral chemogenetic activation of GABAergic neurons in the LH promotes spillage and gnawing behavior. **A)** Schematic of viral infection and representative immunofluorescence images (right panels) showing expression of red fluorescent protein (mCherry, red) and nuclear labeling with 4ʹ,6-diamidino-2ʹ-phenylindole (DAPI, blue) in neuronal somata within LH of Vgat-cre mice. The white scale bar in the lower right corner represents 10 *μ*m. Cumulative feeding time in **B)** fed (control group, Ctrl) and **C)** fasted (Sal) mice following intraperitoneal administration of saline. **D)** Feeding behavior in fed mice following intraperitoneal injection of Clozapine-N-oxide (CNO), a ligand for hM3D(Gq). In the feeding ethograms, pink lines represent HFD pellet consumption, yellow lines indicate Chow pellet consumption, and black lines denote gnawing events. In cumulative feeding time plots, solid lines show group averages (n = 4 mice), and shaded areas represent the ± SEM. **E)** Average IBIs and feeding rate were calculated in 10-minute bins across the session (top and middle panels, respectively) (two-way ANOVA, time effect: F_(11,108)_ = 4.942, *p* < 0.0001, Treatment effect: F_(2,108)_ = 15.83, *p* < 0.0001, and interaction: F_(22,108)_ = 2.262, *p* = 0.0030; *post hoc* analysis: Fed Saline vs. Fed CNO time bins = 0.17 h, 0.33 h, 0.5 h, All *p’s* < 0.05; Fasted Saline vs. Fed CNO time bins = 0.17 h, 0.33 h, 0. 5 h, 0.83 h; All *p’s* < 0.05). In the feeding rate measure, red dots indicate a significant difference between groups. * Indicate statistically significant differences between Ctrl vs CNO (Saline/Fed vs CNO/Fed), and a # symbol denotes significant differences between Sal vs CNO (Saline/Fasted vs CNO/Fed). Bottom panel: Scatter plots display correlations between feeding time and food intake with or without spillage (left and right panels, respectively). Red lines indicate the correlation considering all groups (No spillage R² = 0.32, *p* = 0.5782; Spillage R² = 0.79, *p* = 0.0001), while blue lines denote the correlation excluding the CNO group (No-Spillage R² = 0.55, *p* = 0.0354; Spillage R² = 0.51, *p* = 0.0462). **F)** Quantitative analysis of feeding behavior, including total food intake, spillage, gnawing bout size, number of feeding bouts, bout size, latency to the first bite, IBIs, and intake of a 10% sucrose solution. Data are shown as mean ± SEM. \**p* < 0.05, \*\**p* < 0.01, \*\*\**p* < 0.001. Unilateral chemogenetic DREADD activation of LH GABAergic neurons (n=3 mice) produced similar results (**Supplementary** Fig. 5**-1**).

Analysis of the IBI revealed that Saline/Fasted mice initially exhibited short IBIs, which lengthened as they approached satiety. In contrast, CNO-treated fed mice (CNO/Fed) maintained short IBIs for over an hour (**Figure 5E**, top panel). Similarly, the feeding rate of CNO/Fed mice was sustained above the Saline/Fasted group for up to 50 minutes, indicating that chemogenetic activation of LH GABAergic neurons promotes an intense period of continuous consummatory behavior (**Figure 5E**, middle panel). Interestingly, this intense feeding episode did not directly correspond to increased food consumption. The correlation between feeding time and food intake was weak (R² = 0.32, red line, **Figure 5E**, bottom-left panel) when the CNO/Fed group was included in the analysis. However, this correlation increased dramatically (R^2^ = 0.79, red line) when food spillage was considered (**Figure 5E**, bottom-right panel), suggesting that CNO-treated fed mice were not consuming all the food. A quantitative analysis of feeding microstructure revealed that activating LH GABAergic neurons promoted feeding-related motor patterns without increasing overall food ingestion. Although total food intake was similar across all groups, CNO-treated mice exhibited significantly more food spillage (one-way ANOVA, *F*(2,9) = 48.72, *p* = < 0.0001; *post hoc* analysis: Fed saline vs Fed CNO *p* = < 0.0001, Fasted saline vs Fed CNO *p* = < 0.0001; **Figure 5F**) and gnawing (one-way ANOVA, *F*(2,9) = 19.42, *p* = 0.0005; *post hoc* analysis: Fed saline vs Fed CNO *p* = 0.0004, Fasted saline vs Fed CNO *p* = 0.0005; **Figure 5F**). Consequently, these mice spent significantly more time engaged in feeding behaviors (one-way ANOVA, *F* (2,9) = 7.333, *p* = 0.0129; *post hoc* analysis: Fed saline vs Fed CNO *p* = 0.0049, Fasted saline vs Fed CNO *p* = 0.0238; **Figure 5F**), despite being satiated. This was driven by a change in feeding frequency, as CNO-treated fed mice initiated significantly more feeding bouts (one-way ANOVA, F(2,9) = 3.631, *p* = 0.0698; *post hoc* analysis: Fed Saline vs. Fed CNO, *p* = 0.0367) and had shorter IBIs (one-way ANOVA, F(2,9) = 3.346, *p* = 0.0820; *post hoc* analysis: Fed Saline vs. Fed CNO, *p* = 0.0294). In contrast, the size of each bout (one-way ANOVA, F(2,9) = 1.133, *p* = 0.3641) and the latency to the first bite were unaffected (one-way ANOVA, F(2,9) = 0.107, *p* = 0.8992).

Given that these LH GABAergic neurons have been implicated in reward processing (Gordon-Fennell et al., 2025), we next tested whether their activation would specifically increase consumption of highly palatable food. In agreement with previous findings (Garcia et al., 2021; Ha et al., 2024; Jennings et al., 2015; Navarro et al., 2016), CNO administration prompted a significant increase in liquid sugar intake compared to control groups (one-way ANOVA, F(2,9) = 9.754, *p* = 0.0056; *post hoc* analysis: Fed Saline vs. Fed CNO, *p* = 0.0025, Fasted saline vs Fed CNO *p* = 0.0081; **Figure 5F**). Together, these results suggest that while activating this neuronal population drives a robust motivation to eat, this drive is most effectively translated into consumption when the palatable food is liquid sucrose.

Notably, unilateral DREADD infections in other naïve n=3 Vgat-cre mice yielded results comparable to bilateral infections. While the effect size was slightly reduced with unilateral administration, the difference between the two viral delivery methods was not statistically significant (**Supplementary Fig. 5-1**).

In summary, activating LH GABAergic neurons promoted a general increase in consummatory behaviors—including liquid sucrose intake, feeding time, spillage, and gnawing—without increasing solid food ingestion. These results demonstrate that the Crunchometer can successfully differentiate actual food consumption from other neuronally induced oromotor behaviors, highlighting its value in dissecting complex feeding-related circuits and describing complex non-physiological feeding dynamics. This aligns with previous findings that these neurons increase consummatory behavior toward any available stimulus, including caloric (e.g., food, sucrose), non-caloric (e.g., saccharin), and non-nutritive items (e.g., wood, cork) (Garcia et al., 2021; Jennings et al., 2015; Navarro et al., 2016; Nieh et al., 2015).

### The Crunchometer can be utilized in electrophysiological experiments to align neuronal responses to feeding behavior

After demonstrating that the Crunchometer enables us to capture and quantify hunger and satiety states, as well as how these feeding patterns are disturbed by pharmacological appetite suppressants and chemogenetic manipulations, we employed the Crunchometer to uncover the neuronal responses of the LH associated with feeding behavior at a millisecond timescale (Garcia et al., 2021). We implanted an OptoDrive electrode array in the LH of two mice to record extracellular activity (Caballero-Ruiz et al., 2025). To synchronize signals, a pulse generator system was implemented (**Supplementary Fig. 6-1**), whose output was recorded both through the OpenEphys electrophysiology system’s analog input and via an LED that blinked on and off (1 Hz, 55% duty cycle), captured frame by frame in the video. This setup allowed alignment of the analog signal, LED blinking, and raw audio using the onset of the first pulse as a common reference (**Figure 6A-B**).

**Figure 6.**
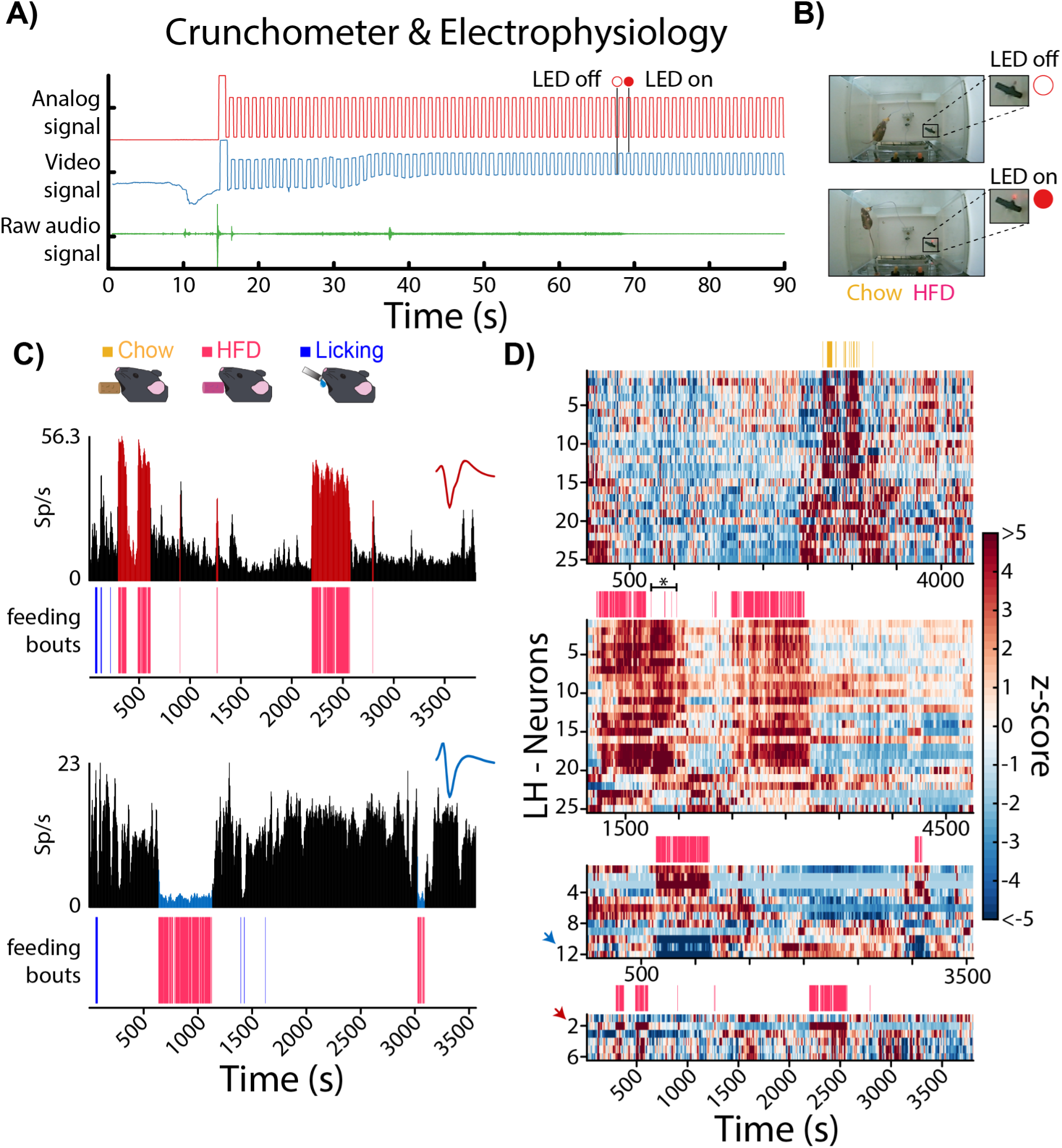
The Cruchometer and electrophysiological responses of LH neurons in freely behaving mice. **A)** To record electrophysiological responses with the Crunchometer, a pulse generator system synchronizes the Crunchometer data with neuronal recordings. Video and audio signals captured by the Crunchometer are aligned with the electrophysiological recordings using a pulse generator, whose output is simultaneously recorded through the electrophysiology analog input (red line) and used to blink an LED (blue line), which is captured frame by frame by the camera during the session. **B)** Example of LED blinking used to synchronize video and audio signals with electrophysiological recordings. Frames showing the LED off (top) and on (bottom) were subtracted from the video to generate a signal resembling the analog input; these frames correspond in time to the horizontal black lines in panel **A**. In **Supplementary** Fig. 6**-1,** there are more details of how to build a 1 Hz pulse generator to blink an LED. **C)** Representative activity of LH neurons during feeding behavior. The firing rates of individual LH neurons were calculated (top plots) and aligned to feeding bouts (bottom plots). The top panel exhibits a neuron that increased its firing rate during feeding bouts (red bars), whereas the bottom panel displays a neuron that decreased its firing rate during the feeding bouts (blue bars). Feeding bouts for Chow and HFD pellets are marked by yellow and pink squares, respectively; licking behavior is indicated by blue squares. The inset depicts the action potential waveform in two representative neurons. **D)** Population activity from four LH neuronal recording sessions. Top plots display feeding bouts in yellow and pink for Chow and HFD pellets, respectively; bottom plots exhibit the z-scored neuronal population activity of the LH across sessions. Red and blue arrows highlight the representative neurons in panel **C** that were selectively activated or inhibited during food intake, respectively.

We recorded a total of 68 LH neurons from two freely behaving mice (two sessions per mouse) during the feeding behavior. LH neurons exhibited either increases or decreases in firing rates during feeding and remained modulated specifically throughout sequences of closely timed feeding bouts, defined as a meal (a collection of feeding bouts punctuated by pauses larger than 2.5 min; **Figure 6C**). Interestingly, this analysis revealed for the first time that individual feeding bouts *per se* do not modulate LH neurons; instead, they exhibited neuronal modulation that spanned the entire meal. Accordingly, we named these LH neurons meal-related neurons. Previously, we reported similar meal-related neurons associated with Ensure intake in the nucleus accumbens shell (Tellez et al., 2012). Then, we identified that 63.24% of neurons were activated (n = 43), 22.06% were inhibited (n = 15), and 14.70% exhibited no modulation by feeding behavior (n = 10; **Figure 6D**), suggesting that LH activity mainly increased during the consumption of solid food. The high-temporal resolution of audio signals underscored the reliability of the Crunchometer in precisely aligning neuronal activity with feeding bouts. Impressively, LH responses were robust enough to detect feeding behavior even under a food hoarding event (see asterisk and horizontal line in **Figure 6D**), when the Crunchometer did not detect bite sounds, such as when the mouse cut off a large chunk of an HFD pellet and chewed it away from the microphone, reducing the accuracy of the sound-based detection (see **Video 4**). Thus, our data demonstrates that the act of chewing food robustly modulates LH neurons, even when the mice are eating and moving around the box simultaneously.

### Event-Locked Alignment of Calcium Recordings to Acoustically Detected Feeding Bouts via the Crunchometer

To assess the utility of the Crunchometer for uncovering neuronal correlates of feeding behavior, we conducted pilot experiments combining microendoscopic calcium imaging with behavioral monitoring in freely moving mice. Animals expressed GCaMP7s in either GABAergic (Vgat-Cre; n = 2 mice, 3 sessions) or glutamatergic (Vglut2-Cre; n = 3 mice, 6 sessions) neurons of the LH. During 30-minute sessions, mice freely fed in the Crunchometer arena, which provided simultaneous access to Chow pellets, HFD pellets, and a licking port delivering water or sucrose solution (**Figures 7A** and **8A**; right panels). Neurons were identified, and their calcium signals were extracted and normalized to z-scores. Neuronal activity was then aligned to feeding bouts and licking events detected by the Crunchometer and the lickometer contact sensor, respectively (**Figure 7B**, top and bottom). In Vgat mice, calcium imaging revealed heterogeneous activity patterns within the LH GABAergic population during feeding bouts, with subsets of neurons showing either increased or decreased calcium transients (**Figure 7C**). Neurons inhibited during feeding likely reflect local inhibitory interactions among LH GABAergic cells, a dynamic reminiscent of the ONsemble/OFFsemble organization described in cortical circuits, wherein activity in one neural ensemble is mirrored by suppression in a complementary ensemble (Pérez-Ortega et al., 2024). Consistent with this framework, neurons 6, 7, and 8 in the right-panel experiment (**Figure 7C**) define a feeding ONsemble, while neurons 3 and 14 display reciprocally suppressed activity characteristic of an OFFsemble. Together, these findings indicate that food consumption engages structurally opposed ensembles within the LH GABAergic circuit, suggesting a push-pull architecture that may coordinate the magnitude or duration of feeding responses. To formally quantify the relationship between individual neuronal activity and ingestive behavior, we computed receiver operating characteristic (ROC) curves assessing how reliably single-unit activity predicted the occurrence of feeding or licking events (**Figure 7D**). Neurons whose decoding performance departed significantly from chance (p < 0.05, shuffle test) were classified as event-increasing or event-decreasing. This analysis identified neurons whose activity tracked feeding or licking with varying predictive strength, spanning a continuum from weakly to strongly modulated units (**Figure 7E**). We observed a similar proportion of GABAergic neurons modulated by solid food feeding (n=32 out of 79; 41%) and liquid licking (n=24 out of 52; 46%). When feeding and licking occurred within the same session, plotting each neuron’s feeding versus licking AUC values revealed three functional subpopulations: neurons selectively tuned to feeding, neurons selectively tuned to licking, and neurons differentially modulated across both behaviors (**Figure 7F**). Notably, the subset of LH GABAergic neurons responsive to both solid and liquid food (n=28 out of 52; 54%) displayed opposite-sign modulation, increasing during one behavior while decreasing during the other, suggesting that these cells do not simply encode ingestive behavior in general, but rather discriminate between solid and liquid food types.

**Figure 7.**
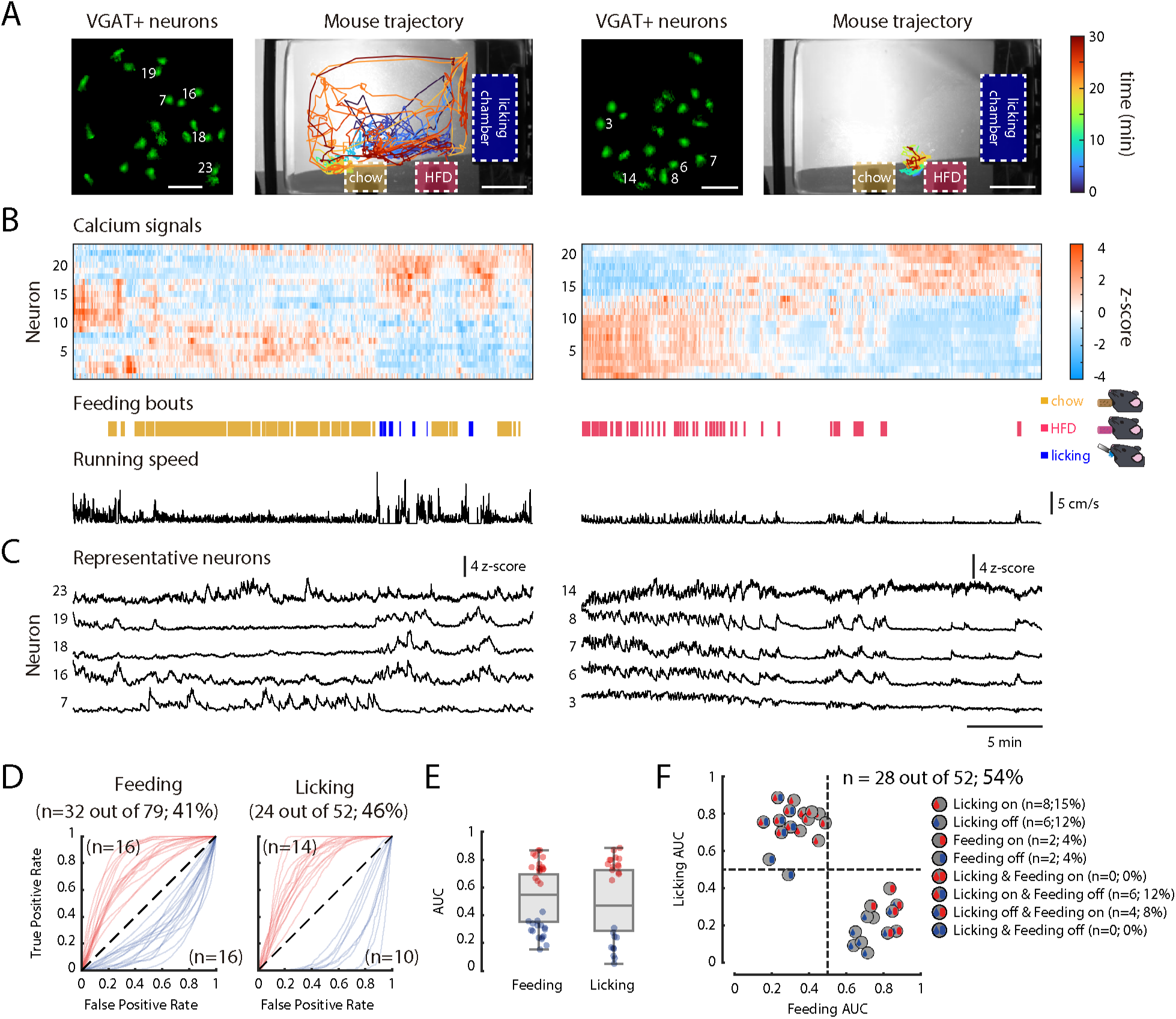
Microendoscope calcium imaging of LH GABAergic neurons during liquid and solid food intake. **A)** Two pilot experiments are shown (left and right panels). Left: GCaMP7s fluorescence images displaying ROIs of individual GABAergic neurons in the LH. Scale bar = 20 µm. Right: Mouse trajectories over a 30-minute session with access to Chow pellets, HFD pellets, and a licking chamber delivering water or sucrose solutions. The color gradient indicates time progression (blue to red). Dashed boxes indicate the locations of Chow, HFD, and licking zones. Scale bar = 5 cm. **B)** Top: z-scored calcium traces from all detected neurons for each experiment, sorted by their similarity in activity patterns (Coss et al., 2022; Pérez-Ortega et al., 2024). Bottom: Feeding and licking bouts automatically detected by the Crunchometer, color-coded by food type Chow (yellow), HFD (pink), and licking (blue). Running speed over time, estimated from video tracking. **C)** Calcium traces from representative GABAergic neurons from each experiment. Neurons display diverse activity profiles, including activation and suppression during feeding bouts. In the left panel, neuron 7 exhibited more activity during feeding but no activity during licking sucrose; in contrast, neurons 16, 18, and 19 mainly responded to liquid sucrose but not to solid food. Neuron 23 exhibited activity during both licking and solid food intake, suggesting that it may participate in processing both types of consummatory behaviors. In the panel on the right, during feeding periods with solid food pellets, neurons 6, 7, and 8 become active, while neuron 3 is inhibited, exhibiting a mirror-like pattern, and neuron 14 shows rebound disinhibition. In this session, the mouse did not lick for sucrose. **D)** Receiver operating characteristic (ROC) curves showing the ability of individual neurons to predict feeding (left) or licking (right) events. Each curve corresponds to a neuron whose decoding performance is significantly different from chance (p < 0.05, shuffle test based on temporal shifting of neuronal activity). Only significant neurons are shown. Curves with AUC > 0.5 are shown in red and indicate neurons whose activity increases during the behavioral event, whereas curves with AUC < 0.5 are shown in blue and indicate neurons whose activity decreases during the event. In LH GABAergic neurons, a similar percentage of neurons exhibited significant AUC predictive activity associated with solid-food feeding (n=32/79; 41%) and licking (n = 24/52; 46%) (chi-square (with Yates correction): χ²(1) = 0.21, *p* = 0.646). **E)** Boxplots showing the distribution of AUC values for neurons significantly predicting feeding (left) or licking (right). Only neurons that reached statistical significance in the shuffle test (same neurons shown in **D)** are included. **F)** Scatter plot comparing feeding AUC versus licking AUC for neurons recorded in sessions where both behaviors occurred. Each dot represents one neuron. Dashed lines indicate chance-level decoding (AUC = 0.5) and divide the plot into quadrants corresponding to neurons that significantly predict feeding, licking, or both behaviors. Icons indicate the type of significant response exhibited by each neuron (see legend). The number of significant neurons responsive to liquid and/or solid food relative to the total recorded population is shown in the plot (n=28 out of 52; ∼54%), and the percentage of neurons in each response category is summarized in the legend. A subset of LH GABAergic neurons exhibited significant ROC-based discriminability between solid-food feeding and licking, either through selective responses or through opposite-sign modulation of activity across the two behaviors.

**Figure 8.**
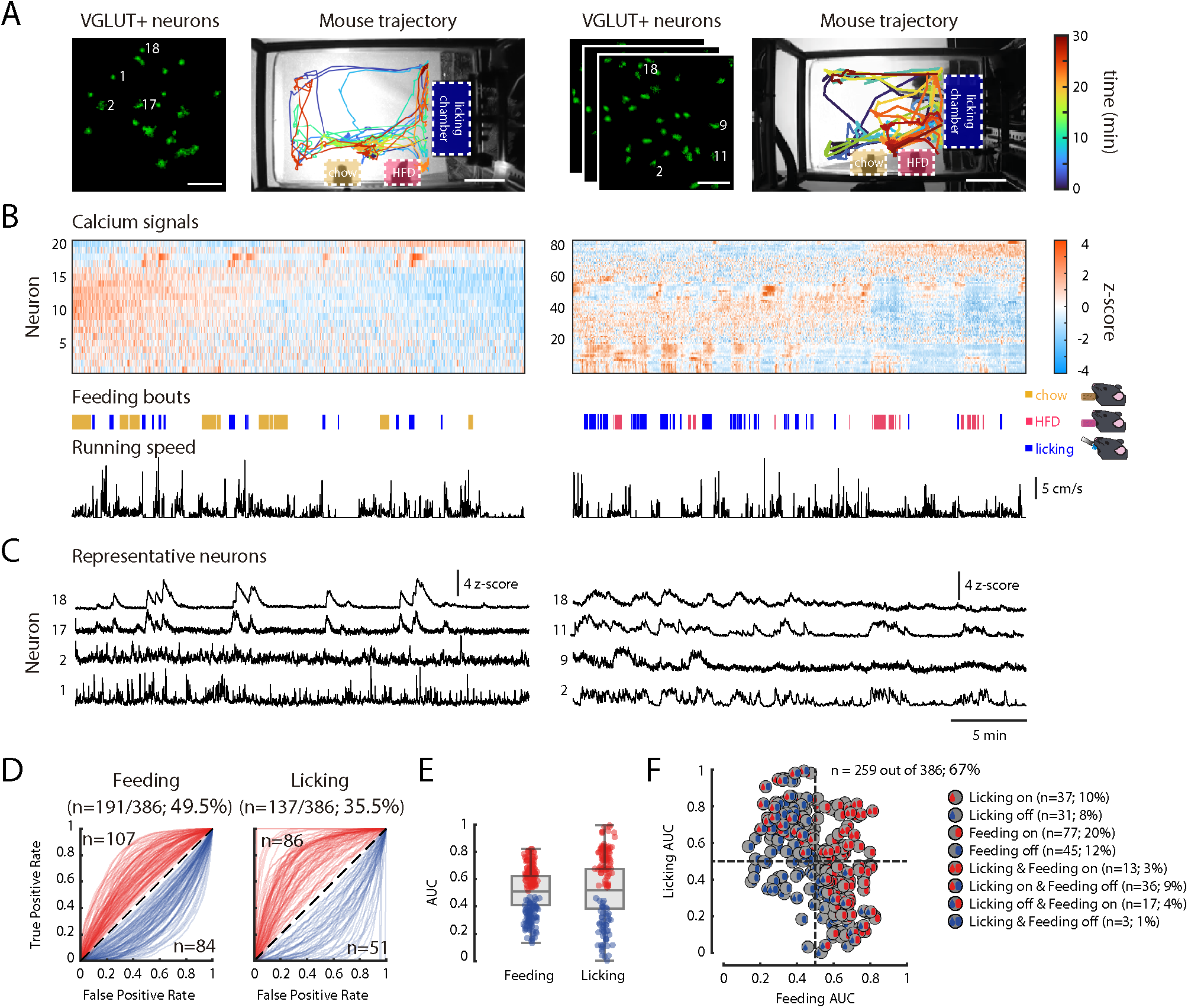
LH glutamatergic neurons differentially encode solid and liquid ingestive behaviors, with a bias toward solid food consumption. **A)** Two representative experiments are shown (left and right panels). Left: GCaMP7s fluorescence image showing ROIs of individual glutamatergic neurons in the lateral hypothalamus. The experiment on the right was performed by recording 3 planes, which allowed us to obtain a larger number of neurons. Scale bar = 20 µm. Right: Mouse trajectories during 30-minute sessions with access to Chow pellets, HFD pellets, and a licking sipper (with a metal floor for a contact lickometer) delivering water or sucrose solutions. Colors represent time progression (blue to red). Dashed boxes indicate the locations of Chow (yellow ticks), HFD (pink), and licking (blue) zones. Scale bar = 5 cm. **B)** Top: z-scored calcium traces from all detected glutamatergic neurons in each experiment. Bottom: Automatically detected feeding and licking bouts as in Figure 7. Mouse’s running speed traces from both experiments. **C)** Example calcium activity traces from glutamatergic neurons illustrating different response profiles across solid and liquid ingestive behaviors. In the experiment on the left, only two neurons (17 and 18) exhibited activity tightly coupled to licking for liquid sucrose but remained unresponsive during the intake of solid food.

In Vglut2-Cre mice, LH glutamatergic neurons similarly displayed heterogeneous response profiles during ingestive behaviors. Example calcium traces illustrate neurons responsive to both feeding and licking (right panel: neurons 2 and 11), neurons selectively tuned to solid food consumption (right panel: neuron 9), and neurons selectively modulated during licking but not during feeding (left panel: neurons 17 and 18; right panel: neuron 18; **Figure 8C**). Applying the same ROC-based decoding framework identified distinct subsets of glutamatergic neurons that were significantly tuned to feeding, licking, or both behaviors (**Figures 8D–F**). Furthermore, among all responsive LH glutamatergic neurons (n=259 out of 386; 67%), a greater proportion was modulated by solid food consumption (49.5%) than by licking (35.5%), indicating that solid food may engage the LH glutamatergic population more broadly (chi-square (with Yates correction): χ²(1) = 14.89, *p* < 0.001). In contrast, this bias is not observed in the GABAergic population, where feeding and licking-modulated neurons were found in similar proportions, suggesting a differential functional organization between LH inhibitory and excitatory circuits.

A key distinction between LH GABAergic and glutamatergic populations further emerged from the AUC scatter plots: unlike GABAergic neurons, glutamatergic neurons exhibited clustering along the diagonal quadrants, indicating concordant modulation (neurons activated during both behaviors or inhibited during both) rather than the opponent-sign, off-diagonal clustering predominant in the GABAergic population (**Figure 8F**). Together, these results demonstrate that LH glutamatergic neurons, like their GABAergic counterparts, are heterogeneously recruited during ingestive behavior and can discriminate between solid food consumption and liquid licking at the single-neuron level.

In the experiment on the right, neurons 2 and 11 exhibited activity associated with both feeding and licking events; neuron 9 showed activity selectively associated with solid food intake, whereas neuron 18 was selectively modulated during liquid consumption. D) ROC curves showing the ability of individual neurons to discriminate feeding (left) or licking (right) events. Each curve corresponds to a neuron whose decoding performance is significantly different from chance (p < 0.05, shuffle test based on temporal shifting of neuronal activity). Only significant neurons are shown. In LH glutamatergic neurons, a greater percentage of neurons exhibited significant AUC predictive activity associated with solid-food feeding (n=191/386; 49.5%), versus those modulated by licking (n = 137/386; 35.5%) (chi-square (with Yates correction): χ²(1) = 14.89, *p* < 0.001). **E)** Boxplots showing the distribution of AUC values for neurons significantly predicting feeding (left) or licking (right). Only neurons that reached statistical significance in the shuffle test (same neurons shown in **D**) are included. **F)** Scatter plot comparing feeding AUC versus licking AUC for neurons recorded in sessions where both behaviors occurred. Each dot represents one neuron. Dashed lines indicate chance-level decoding (AUC = 0.5) and divide the plot into quadrants corresponding to neurons that significantly predict feeding, licking, or both behaviors. Icons indicate the type of significant response exhibited by each neuron (see legend). The number of significant neurons modulated by solid and/or liquid food relative to the total recorded population is shown in the plot (n=259 out of 386; 67%), and the percentage of neurons in each response category is summarized in the legend. Both On and Off-diagonal responses were observed.

## Discussion

This study demonstrates that analyzing bite sounds provides a novel method for characterizing feeding behavior, offering unprecedented temporal resolution and precision for measuring its microstructure with solid foods. This improved temporal resolution for detecting the onset and offset of feeding bouts, based on bite sounds, paves the way for unveiling neuronal modulation phase-locked to feeding, thereby accelerating the discovery of the neural circuits controlling appetite. We propose the Crunchometer as a valuable, low-cost, open-source tool that can be easily implemented in any standard laboratory. To facilitate its adoption, we provide accompanying software along with detailed setup and installation instructions (see (Arroyo et al., 2025)). We also provided a <u>benchmark</u> database (comprising six 2-hour-long recordings of hungry mice, used in **Figure 2** to validate the Crunchometer for further refinement by the scientific community; see the Open Science Frame database (Arroyo et al., 2025).

Our recent research using the Crunchometer (previously named Crunch Master (Liu et al., 2025)) has revealed that the gut can ’taste’ microbial patterns of bacteria, playing a key role in regulating feeding behavior. We discovered that specialized sensory cells in the colon, known as neuropods, utilize a receptor (TLR5) to detect flagellin, a protein produced by certain microbes. This detection triggers the release of the appetite-suppressing hormone PYY. In mice, this signal reduced food intake and prevented obesity, revealing a new sense to detect intestinal biota named the “neurobiotic sense” (Liu et al., 2025). The device’s precision in tracking the consumption of an individual Chow pellet was essential for this detailed microstructural analysis. In this work, we extend the capabilities of the Crunchometer (Liu et al., 2025). By enabling the simultaneous monitoring of two different solid foods (standard Chow and a high-fat diet) alongside a liquid option (sugar water), the system can now be used to assess food preference in addition to the fine-grained analysis of feeding microstructure.

### Microstructure of feeding under different fed and fasted energy states

This study demonstrates that the Crunchometer system accurately differentiates the meal patterns of fed and fasted wildtype mice by quantifying their feeding microstructure. Fasted mice exhibited shorter latencies to eat, higher initial feeding rates, and more frequent bouts, resulting in a doubling of their cumulative feeding time and food intake compared to their satiated counterparts. Furthermore, the initial increase in feeding rate followed by a progressive lengthening of the IBI in fasted mice illustrated the transition to satiety. This ability to resolve significant differences between fed and fasted energy states validates the Crunchometer as a sensitive tool for investigating the neurobiological and pharmacological regulation of appetite (Shrivastava et al., 2025; Wang et al., 2024).

### Semaglutide: more than just appetite suppression, it reshapes fatty, energy-dense food preferences

The Crunchometer analysis revealed rapid and robust appetite-suppressant effects of acute semaglutide administration, consistent with the idea that this GLP-1 agonist induces a robust satiety sensation (Drucker, 2025). More striking was the reduction in preference for the HFD observed on the administration day, which was maintained until the next day (post-semaglutide; **Figure 4**, see **Video 2**). Future studies should use the Crunchometer to characterize changes in HFD/chow preference during 24-h monitoring under chronic semaglutide treatment. This result aligns with observations in humans, in which semaglutide was also associated with reduced hunger and food cravings, improved control over eating, and, more importantly, a lower preference for fatty, energy-dense, non-sweet foods (Blundell et al., 2017).

### Chemogenetics activation of LH GABAergic neurons triggers a stress-induced eating pattern

Activating GABAergic neurons in LH via chemogenetics induces a striking behavioral phenotype. Instead of normal, homeostatic eating, this stimulation triggers a frantic, stress-like pattern of consumption, characterized by a significant surge in biting and gnawing directed at the food source (**Video 3**). The Cruchometer and the Threshold method could successfully capture this frenetic feeding pattern with unprecedented precision. This feeding pattern involved hypertrophic mastication and gnawing directed at all proximal stimuli (palatable, non-palatable, or inedible), resulting in substantial food spillage and increased sucrose liquid intake. In agreement with previous findings (Garcia et al., 2021; Ha et al., 2024; Jennings et al., 2015; Navarro et al., 2016).

### Beyond the lick, a new world of possibilities for the neuronal correlates of eating solid food: Meal-related neurons in the LH

Extracellular multichannel recordings and calcium imaging were easily synchronized with the Crunchometer, allowing for alignment of neuronal responses to the onset and offset of a feeding bout with unprecedented resolution. The study of neuronal correlates of eating solid food has been rarely investigated, see (Pilato et al., 2024; Yamamoto et al., 1989), for exceptions. Here, we found that LH neurons exhibit robust modulation during the consumption of solid foods. We named these responses LH meal-related neurons, in agreement with the well-established role of LH on ingestive behavior. Unexpectedly. Our pilot data (see below for calcium imaging) suggest that licking vs eating solid food may rely upon distinct neuronal circuits, in agreement with Dilorenzo’s finding in rostral NTS neurons, in which they found little correspondence between liquid taste-evoked responses and those evoked by eating solid food. Pat Dilorenzo wrote, “What is striking is that we see no correspondence whatsoever, not even a weak one(O’Connell et al., 2025).” Here we also observed a partial separation of neurons responding to solid food and liquid sucrose (more of this below). The fact that licking vs. eating solid food seems to recruit distinct neuronal ensembles also opens the possibility of labeling them using TRAP or Tet-tagging systems (Guenthner et al., 2013; Zhang et al., 2015) to further test their specific contribution to feeding.

### LH GABAergic neurons discriminate between solid and liquid food either through selective responses or through opposite-sign modulation of activity across the two behaviors

Calcium imaging in Vgat::GCaMP7s mice revealed that LH GABAergic neurons are heterogeneously modulated during ingestive behavior, with distinct subpopulations preferentially tuned to solid food consumption, liquid licking, or both. Among neurons responsive to both, GABAergic cells encoded solid and liquid ingestion mainly through opposite-sign modulation (whereby the same cell increases its activity during one ingestive event while decreasing it during the other), suggesting that individual LH GABAergic neurons may act as comparators that weigh sensory or motivational signals associated with distinct food types, rather than simply reporting the presence of liquid or solid food. Moreover, the presence of a rich proportion of licking-modulated neurons within the responsive GABAergic population further suggests that liquid reward reliably drives LH GABAergic activity. Together, both the opponent-sign discrimination could reflect differences in the orosensory, textural, or postingestive properties of solid versus liquid stimuli, and may be relevant to the well-documented capacity of animals to independently regulate solid and liquid intake. Consistent with this interpretation, LH neurotensin-expressing neurons (a molecularly defined GABAergic subpopulation) selectively promote fluid intake while modestly suppressing (or not affecting) solid food consumption (Kurt et al., 2019). This is further consistent with the well-documented capacity of LH GABAergic neuron activation to elicit orolingual stereotypies (Garcia et al., 2021; Jennings et al., 2015; Navarro et al., 2016; Nieh et al., 2015). Although based on a small sample (n = 2 mice, 3 sessions, and 79 neurons), these preliminary observations demonstrate the utility of combining the Crunchometer with calcium imaging to reveal behaviorally relevant neural dynamics. Future studies with larger cohorts will be needed to determine whether opponent-sign modulation reflects functionally distinct cell-type subpopulations or state-dependent switches within individual neurons.

### LH glutamatergic neurons also differentially encode solid and liquid ingestive behaviors, with a bias toward solid food consumption

In recordings from VGlut2::GCaMP mice, glutamatergic neurons also exhibited heterogeneous response profiles across ingestive behaviors. Example neurons showed activity associated with both feeding and licking, selective responses to feeding events, or modulation restricted to licking behavior. ROC-based decoding analysis revealed subsets of neurons whose activity significantly predicted feeding, licking, or both behaviors. While prior work has established that LH glutamatergic neurons respond to liquid sucrose (Gordon-Fennell et al., 2025; Rossi et al., 2019), the present results extend this picture: a larger fraction of glutamatergic neurons displayed significant predictive activity for solid food consumption than for licking, demonstrating that this population also actively participates in encoding solid-food intake under naturalistic, freely moving conditions. While these observations are based on a limited number of recordings (n= 3 mice, 6 sessions, and 386 neurons), they highlight the value of the Crunchometer and suggest that future studies with expanded datasets will be crucial to determine how glutamatergic LH neurons contribute to ingestive control under different behavioral contexts, including those related to reward or aversion (Gordon-Fennell and Stuber, 2021).

### Limitations and Future Directions

While the Crunchometer provides accurate temporal detection of bites and feeding microstructure, estimating absolute food mass consumed from bite-related acoustic signals shows considerable variability across trials and subjects. This limitation most likely arises from individual differences in gnawing patterns, food fragmentation, and hoarding behavior. Accordingly, the Crunchometer is best suited for analyses of feeding dynamics and behavioral microstructure, whereas studies requiring precise quantification of ingested mass should complement the system with direct gravimetric measurements (for example, real-time weighing of feeders (Rathod and Di Fulvio, 2021)). A methodological consideration is that chemogenetic activation via DREADDs imposes a sustained, supra-physiological drive that most likely does not reproduce the temporal structure of endogenous LHA GABAergic activity during feeding; optogenetic manipulations share analogous limitations (see *optoception*; (Luis-Islas et al., 2022)). Our findings, therefore, establish that activation of this neuronal population is sufficient to produce uncontrolled feeding and gnawing, without implying that its endogenous firing encodes them in the same manner. Furthermore, our current SVM model reliably distinguishes between bites, non-bites (silence), and artifacts; however, it does not accurately differentiate between biting and gnawing, likely due to highly similar acoustic profiles recorded by our microphone. As a result, gnawing periods were manually reviewed. Nevertheless, our current SVM model for the Crunchometer provides a fully automated and scalable method for identifying bite events from audio recordings, with minimal human intervention limited to initial model training and fewer artifact detections. Future iterations of the classifier could incorporate finer spectral features or unsupervised pre-clustering to disambiguate subtle behaviors. Another limitation of our trained SVM model is that it is expected to generalize poorly across different Crunchometer setups (not shown). Consequently, retraining a new model is likely required for each laboratory or for any setup that uses a microphone different from the one standardized here. In this regard, the Threshold method is more reliable and easier to tune for each setup. That said, to our knowledge, this is the first application of machine learning to detect mouse feeding microstructure acoustically across varied experimental paradigms. Other impressive deep-learning methods have been previously developed, but they were designed to classify and sort distinct vocalization types, not to detect feeding bouts (Coffey et al., 2019). For a more comprehensive behavioral analysis, the Crunchometer’s feeding ethogram can be supplemented (but at a higher computational cost) by DeepEthogram a video software (Bohnslav et al., 2021), to identify other complex activities, such as grooming. Closed-loop optogenetic experiments were not tested here, but they could also be easily implemented by detecting bites in real time and sending TTL pulses to control laser activation. For these closed-loop experiments, our SVM model’s predictions are ideal for rapidly detecting feeding bouts with a low artifact rate. Finally, long-term recordings: the size of the video files, though not the audio files, makes continuous 24-hour recording more challenging to process (but not impossible) due to the large storage space required. While technically possible (in fact, we have now optimized the Python version and the standalone MacOS Crunchometer software to open and process 24 h .mkv videos), another approach for long-term studies would be to adapt the Crunchometer to an event-triggered recording, limited to feeding events (saving video and sound snippets) in real time. We anticipate that future versions of the Crunchometer will incorporate closed-loop capability, contributing a powerful new behavioral tool towards achieving the long-term dream of Curt P. Richter. We anticipate the Crunchometer will allow the neuroscience field to “freely” move forward from head-fixed approaches and further enrich the new era of the “behavioristic study of the activity of rodents (Richter, 1922).”

## Conclusion

The Crunchometer offers a transformative and accessible tool for high-resolution studies of solid food consumption, paving the way for democratizing the use of these technologies (Marzullo and Gage, 2012). By integrating effortlessly with essential techniques such as *in vivo* electrophysiology and calcium imaging, it has already yielded novel insights into the neural basis of feeding in freely behaving mice. These findings include the discovery that bacterial microbial patterns regulate mice’s feeding behavior (Liu et al., 2025), the specific role of LH GABAergic neurons in feeding, and the distinct neural encoding of solid versus liquid food rewards. As a powerful, ready-to-deploy hardware-and-software solution, this low-cost, open-source technology promises to accelerate the precise dissection of appetite circuits and the development of new therapies for obesity and eating disorders.

## Material and Methods

### Subjects

Adult male and female C57BL/6J mice (20–40 g) were used to study energy states (hunger and satiety) and the anorexigenic effects of semaglutide. Separately, adult male and female Vgat-cre mice (JAX:016962 STOCK Slc32a1tm2(cre)Lowl/J) were used for chemogenetic activation of GABAergic neurons or for electrophysiological recordings. Vgat-cre and Vglut2-cre ( JAX:016963 STOCK Slc17a6tm2(cre)Lowl/J) adult mice were used for calcium imaging recordings. All mice were individually housed in standard laboratory cages under controlled conditions: a 12:12 h light/dark cycle (lights on at 06:00), a temperature of 22 ± 2°C, and a relative humidity of 50 ± 5%. Mice had *ad libitum* access to water and standard Chow diet (LabDiet 5008) unless otherwise stated. No sex differences were observed in any of the experiments; therefore, data from male and female mice were pooled. All behavioral experiments were carried out during the light phase of the cycle. All procedures involving animals were approved by the Institutional Animal Care and Use Committee (IACUC) of CINVESTAV.

### Viral vector

The cre-inducible adeno-associated virus (AAVs) were purchased from Addgene (Watertown, MA, USA). The viral concentration was 4×10¹² viral genomes ml-¹ for AAV8-hSyn-DIO-hM3D(Gq)-mCherry (Addgene #44361). pAG-AAV-syn-FLEX-jGCaMP7s-WPRE (Addgene #104491) virus was used for calcium imaging experiments. Viruses were divided into aliquots and stored at -80 °C until use.

### Stereotaxic surgery

Mice were anesthetized with isoflurane (induction, 5%; maintenance, 1–1.5%; ViP 3000 Matrix). The microinjection needles (29-G) were connected to a 10 μl Hamilton syringe and filled with AAV. For all experiments, the mice were bilaterally injected into the LH (from Bregma (mm): −1.4 AP, ±1.0 ML, and −5.8 DV) with AAV (0.25 μl) at a rate of 0.1 µl min−1 with an additional 5 min for a complete diffusion. Coordinates were taken from Allen’s reference atlas of the mouse brain. Following the suturing of the mouse’s head, a three-week recovery period was allowed to enable healing and transgene expression. For electrophysiology, in two Vgat-cre mice, a custom-made 16-tungsten OptoDrive array was implanted targeting the LH (from Bregma (mm): −1.3 AP, ±1.1 ML, and −5.3 DV) (Caballero-Ruiz et al., 2025). The OptoDrive was fixed to the skull using dental acrylic and anchored with a grounding screw. After surgery, mice were allowed to recover for one week, and ketoprofen (45 mg/kg, i.p.) was administered for three consecutive days to manage postoperative pain. For calcium imaging experiments, two Vgat-cre and three Vglut2-cre mice were used. To label neurons with a genetically encoded calcium indicator, an adeno-associated virus (AAV) encoding GCaMP7s was injected into the LH (from Bregma (mm): at coordinates −1.3 AP, ±1.1 ML, and −5.3 DV). A total volume of 300 nL was delivered at a rate of 30 nL/min, and the injection needle was left in place for an additional 10 minutes to allow for viral diffusion. Following surgery, mice were allowed to recover for three weeks to ensure robust GCaMP7s expression. After the recovery period, a gradient refractive index (GRIN) lens (ProView Integrated Lens, 7.3 mm length, 0.6 mm diameter; Inscopix) was implanted at a final depth of DV –5.1 to – 5.2 mm, approximately 200 µm above the injection site. During implantation, the nVISTA microendoscope system (Inscopix) was temporarily connected and activated to verify both GCaMP7s expression and an appropriate field of view. To minimize tissue compression and accommodate displacement caused by the lens, it was advanced in 300 µm increments, with a 100 µm upward retraction after each step. After lens implantation, mice were allowed to recover for an additional five days before the start of behavioral and calcium imaging experiments. To manage postoperative pain, Ketoprofen (45 mg/kg, i.p.) was administered once daily for three consecutive days following both the viral infection and the lens implantation procedures.

### Drugs

Semaglutide was diluted in 0.9% saline to a final concentration of 0.067 mg/mL. Saline and semaglutide were administered via subcutaneous injection at a dose of 0.123 mg/kg immediately before the start of the behavioral experiment. Clozapine-N-Oxide (CNO) was also prepared in 0.9% saline at a final concentration of 2 mg/mL. Both saline and CNO were administered via intraperitoneal injection at a dose of 3 mg/kg immediately before the start of the behavioral protocol.

### Behavioral protocol

All mice were habituated to the Crunchometer for 2 days before the recording session. Each habituation session lasted 30 minutes, during which two food pellets were placed in the chamber: one standard Chow pellet (LabDiet 5008) and one highly palatable high-fat diet (HFD) pellet (Research Diet, D12451). As a practical note, we recommend allowing the HFD to equilibrate to room temperature before the experiment and pre-exposing mice to a single HFD pellet in their home cage to attenuate neophobia prior to testing. Our study is limited to the acoustic detection of standard Chow and HFD pellets, both of which exhibit a firm, brittle consistency. Consequently, future research should evaluate the fidelity of the Crunchometer in characterizing a broader range of food textures, encompassing varying degrees of hardness and elasticity, as reported by (O’Connell et al., 2025). The pellets were positioned on the chamber walls in the same location described in detail in **Figure 1**. During habituation, the mice were in a fed state and allowed to explore both food types freely (no video recordings were performed). This was done to reduce stress and novelty on the day of the behavioral recording. On the recording day, each Crunchometer session lasted two hours. Throughout this period, mice had *ad libitum* access to both standard Chow and HFD pellets. In addition, a 10% sucrose solution (from Sigma-Aldrich Mexico) was provided as a liquid source during the feeding behavior. All fasted mice, regardless of the experimental group, were food-deprived 18 hours before the session, with only solid food removed. In contrast, fed mice had *ad libitum* access to both food and water. For pharmacological and chemogenetic experiments, drug administration was performed immediately before placing the mice in the recording chamber. At the end of each session, mice were returned to their home cages. Food pellets were weighed before and after each session, and any spillage was collected and weighed to calculate precise intake. For calcium imaging, mice were trained in a behavioral chamber containing five stimuli: three liquid solutions (water, 3% sucrose, and 18% sucrose) and two types of food pellets (standard Chow and HFD). Before the calcium imaging recording session, mice underwent a 3-day habituation period (30 minutes/day). On the day of calcium imaging, mice were food-deprived for 18 hours before the session. Each imaging session lasted 30 minutes, during which mice again had free access to the same liquid and solid stimuli.

### Electrophysiological recordings

Electrophysiological recordings were performed using the OpenEphys Acquisition Board (OpenEphys Production Site) and GUI (Siegle et al., 2017) during feeding behavior in freely moving mice. The neural signals were sampled at 30 KHz and band-pass filtered between 0.5 and 8 KHz. Action potentials exceeding 50 µV were recorded, and putative units were identified online using voltage-time windows. Subsequently, offline spike sorting was implemented using an unsupervised algorithm (Chaure et al., 2018) to isolate single units. Only the timestamps of these sorted units were used for further analysis.

### Immunofluorescence

Mice were initially sedated with isoflurane (5%, ViP 3000 Matrix) for 1 minute and subsequently euthanized using a lethal dose of embutramide and mebezonium iodide (T61) (2 ml/kg). Intracardiac perfusion was performed using phosphate-buffered saline (PBS), followed by fixation with 4% paraformaldehyde in 0.1 M phosphate buffer. The brain was removed and fixed in 4% paraformaldehyde and then stored for three days. Brain tissues were dehydrated by gradually increasing the sucrose concentration (10%, 20%, and 30%) before slicing. Brains were sliced into 40 μm sections using a cryostat (Thermo Scientific HM525). Free-floating sections were incubated with 300 nM 4’, 6-diamidino-2-phenylindole (DAPI, D9542, Sigma-Aldrich) for 1 min. After a final wash with TBST, sections were mounted in Dako fluorescence mounting medium. Immunofluorescence was observed using a LEICA Stellaris 5 confocal microscope.

### Signal processing pipeline of the Crunchometer

In each recording session, we obtained an “.mkv” file containing both video and audio signals. From the video file, the audio was extracted as an “.ogg” file for further analysis using a custom MATLAB function (extractAudio.xml) or in the Crunchometer software (Tab 1 (Processing Tools). 1. Audio Extractor). To analyze the audio, the power spectrum was calculated using a short-time Fourier transform (using MATLAB’s spectrogram function). This produced a matrix in which each column represented a bin of a 1-second time window (with no overlap), and each row corresponded to a specific frequency. We focused on the 500-950 Hz frequency band, which was associated with bite sounds produced by the mice. The bite-related power was computed as follows:

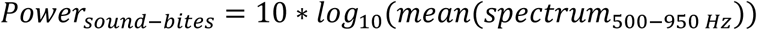

To detect sound bites, we set a threshold based on the distribution of power values across sessions. This threshold was defined as:

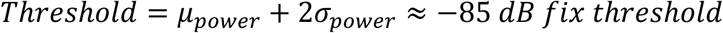

This threshold value was consistent across sessions (corresponding to ∼3 std above the mean), given our standardized experimental conditions. Bins exceeding the threshold were classified as detections and grouped into sequences with less than 5 seconds between detections, referred to as feeding bouts. At a macrostructural level, a meal was defined as a pause between feeding bouts lasting longer than 2.5 minutes. Once the time intervals of these bouts were identified, they were used to generate “video snippets.mp4” video files containing both audio and video around each possible feeding event. For automated sorting, each snippet was placed into a labeled folder (e.g., Artifact, Chow, HFD, Gnawing) based on the mouse’s proximity to either the Chow or HFD pellet locations during the event. If the mouse was not near either pellet source, the detection was classified as an Artifact. Then, all snippets were reviewed to confirm or correct their categorization. Misclassified snippets were manually reassigned to their respective folders.

### Statistical Analysis

Data on cumulative feeding time were analyzed using a two-sample Kolmogorov–Smirnov test (two-tailed) to assess differences in distribution. The one-tailed Mann–Whitney U test for group comparisons, as well as one-way and two-way ANOVA, were applied, where appropriate, to the crunchometer metrics. Fisheŕs LSD post hoc analysis was performed. *α* was always set *p* < 0.05.

### SVM model of the Crunchometer: automatic detection of biting from audio

We developed a supervised classification algorithm based on a multi-class support vector machine (SVM), implemented in MATLAB, to automatically detect biting events from audio recordings. The classifier was trained on data from a single 1-hour session recorded from one mouse. Audio segments were manually annotated into three categories: “bite”, “artifact”, and “no sound”. Recordings were acquired at 44,1 kHz using a mono microphone placed inside the behavioral chamber. Spectrograms were computed using 1-second non-overlapping windows with a frequency resolution of 5.4 Hz (nfft = 2¹³). Each time window was represented as a column vector of spectral features. Pairwise Euclidean distances between these vectors were calculated and used for hierarchical clustering with the Ward linkage method. The resulting clusters were manually assigned to one of the three sound categories (bite, artifact, or no sound) based on an inspection of the spectrogram patterns, associated audio, and synchronized video clips. This labeled dataset was then used to train the multi-class SVM. Remarkably, this model, trained on data from a single mouse, generalized robustly across all mice recorded under similar conditions, without requiring additional retraining. For subsequent recordings (sampled at 44,1 kHz), spectrograms were computed using the same parameters, and each 1-second window was classified as a bite, artifact, or silence using the trained model.

### Definition of feeding bouts

Feeding bouts were defined as sequences of bite events separated by less than 5 seconds. If more than 5 seconds elapsed between bite detections, a new bout was initiated. Feeding bouts were categorized as “Chow” or “HFD” based on video analysis of mouse location and movement (see below). Licking events were identified separately using a lickometer and were not included in feeding bouts.

### Identification of Chow vs. HFD consumption

To identify the type of pellet consumed during each feeding bout, we initially analyzed the spectrogram within the frequency band associated with sound bites. However, no differences were found among pellets based on their acoustic features. Therefore, we examined the video clips corresponding to the detected feeding intervals. Two regions of interest (ROIs), defined by the location of the pellets in the video, were manually defined over the Chow and HFD pellet dispensers. Motion energy was computed for each ROI using the MoussionEnergy algorithm (Pérez-Ortega, 2023a), based on the mean absolute difference between consecutive video frames. If the motion energy within an ROI exceeded a manually set threshold (determined to exclude noise), the bout was classified accordingly: “Chow” if the Chow ROI had greater motion energy, “HFD” if the HFD ROI did. Bouts with low or ambiguous motion were excluded.

### Calcium imaging experiments and analysis

GCaMP7s-expressing Vgat-cre or Vglut2-cre mice were implanted with Inscopix microendoscopes targeting the lateral hypothalamus. Calcium imaging was conducted using the Inscopix nVISTA microendoscope system (Palo Alto, CA, USA). Neural activity was recorded continuously for 30 minutes at a sampling rate of 30 Hz (Inscopix Data Processing Software). To enable synchronization with behavior, a trigger signal was delivered at the onset of the experiment. Behavioral events, including individual licks, were precisely timestamped. Additionally, a low-frequency auditory tone (250 Hz, dB = 67.1) paired with a visible LED flash in the video, both synchronized and recorded, provided a reference point to ensure alignment between the behavioral video and neural datasets. This LED/tone pairing was also timestamped within the nVISTA system (Inscopix), facilitating accurate temporal correlation between calcium dynamics and specific behavioral actions. Raw calcium videos were preprocessed by subtracting a Gaussian-blurred version of each frame (σ ≈ 16 µm) to remove background fluorescence. Rigid-body motion correction (translation only) was then applied. Following the procedure described in (Pérez-Ortega et al., 2024), neuronal regions of interest (ROIs) and ΔF/F₀ fluorescence traces were extracted using the Xsembles2P algorithm (Pérez-Ortega, 2023b). Neurons with a peak signal-to-noise ratio (SNR) below 10 dB were excluded from further analysis. The remaining fluorescence traces were normalized using z-scoring.

### Tracking trajectory and running speed

Mouse position was tracked from grayscale video recordings. A static background was estimated by computing the maximum projection across all frames, resulting in a bright background image. Each frame of the original video was then subtracted from this background, highlighting regions where the mouse was present as darker areas. Foreground segmentation was performed using global thresholding (Otsu’s method), followed by morphological erosion with a disk-shaped structuring element (radius = 2 pixels) to reduce noise. The centroid of the resulting binary mask was used to estimate the mouse’s (x, y) position in each frame. Running speed was calculated as the frame-to-frame displacement, scaled by the video’s sampling rate. Tracking was not possible while the mouse was inside the licking chamber due to occlusion. This was implemented with a homemade MATLAB function Mass_Center_From_MP4.m. In the Crunchometer software on Tab 5 (Trajectory Analysis). Trajectory Analysis can also be used to track the position and velocity of the mouse centroid using a Kalman filter, along with other postural measurements.

### ROC-based decoding of feeding and licking behaviors

To quantify the relationship between neuronal activity and ingestive behaviors, we used receiver operating characteristic (ROC) analysis to test whether calcium signals could predict feeding or licking events. For each neuron, the fluorescence signal was normalized as a z-score and used to predict binary behavioral variables (feeding or licking). A set of 100 thresholds, evenly distributed across the percentile range of the fluorescence signal, was used to compute ROC curves and their corresponding area under the curve (AUC). ROC analyses were performed independently for feeding and licking events. Only neurons showing significant predictive power for at least one behavior were included in the ROC visualizations and AUC summary plots shown in the figures.

### Identification of ON and OFF neurons predicting feeding and licking

To assess statistical significance, we generated 1,000 surrogate datasets by circularly shifting the neuronal activity in time relative to behavioral events and recomputing the ROC curve and AUC for each surrogate. The empirical AUC was then compared to the surrogate distribution. Neurons were considered significant when the empirical AUC differed from the surrogate distribution at p < 0.05. When the empirical AUC exceeded the surrogate values, the neuron was classified as activated (predictive increase in activity; red in figures), whereas when the empirical AUC was lower than the surrogate distribution, the neuron was classified as inhibited (predictive decrease in activity; blue in figures). For sessions in which both feeding and licking occurred, we further compared the selectivity of individual neurons by plotting the AUC obtained for feeding versus the AUC obtained for licking. Based on their significant responses, neurons were assigned to one of eight response categories: Licking on, Licking off, Feeding on, Feeding off, Licking & Feeding on, Licking on & Feeding off, Licking off & Feeding on, and Licking & Feeding off. Only neurons with significant decoding for at least one behavior were included in the ROC and AUC plots. However, response category proportions were calculated relative to the total number of recorded neurons, including those that did not reach significance for either behavior.

## Acknowledgement

We thank Fabiola Hernandez Olvera for invaluable animal care, Mario Gil Moreno for building the OptoDrive. We also thank the Unidad de Imagenología of the Centro de Investigación sobre el Envejecimiento for the image processing, especially to Tzindilu Molina Muñoz.

## Author Contributions

**Conceptualization:** R.G., A.C., and D.V.B. conceived the initial idea for the Crunchometer.

**Methodology:** E.G.-L, B.A., R.G.

**Software:** B.A., A.L., J.P-O., R.G.

**Validation:** E.G.-L., B.A., G.H., A.L., X.D., N.R., R.G.

**Formal analysis:** E.G.-L., B.A., J.P-O., E.H.L, R.G.

**Investigation:** E.G.-L., B.A., X.D., L.A.R.B.

**Resources:** R.G., E.A., E.H.L., M.K., D.V.B.

**Data curation:** E.G.-L., B.A., J.P-O., R.G.

**Writing—original draft:** E.G.-L, B.A., R.G.

**Writing—review & editing:** E.G.-L., B.A., J.P-O., A.L., L.A.R.B., X.D., G.H., A.C., E.A., N.R., E.H.L., M.K., D.V.B., and R.G.

**Visualization:** E.G.-L., B.A., G.H., J.P-O., R.G.

**Supervision:** E.H.L., D.V.B., R.G.

**Project administration:** R.G.

**Funding acquisition:** N.R., M.K., D.V.B., E.H.L., and R.G

## Data and Code Availability

Benchmark and Crunchometer software will be available on OSF (Arroyo et al., 2025) using the following link: https://osf.io/bmkdc/?view_only=913feef710714d2bbd4cfb45960fb7be

The OSF repository contains two folders under **Files**:

**TheCrunchometerV2_Matlab** includes both the classic scripts and an App GUI version, compatible with Windows, macOS (Intel and Apple Silicon), and Linux. This version requires a MATLAB license and FFmpeg and is limited to videos up to 4 hours long.

The **TheCrunchometerV2_Python** folder contains a standalone macOS Silicon executable (TheCrunchometer-2.0.0-macOS-arm64.dmg); this is the recommended version. It requires no MATLAB license or additional dependencies, supports 24-hour video files, and runs optimally on Apple Silicon hardware (e.g., Mac mini M4). To install, double-click the .dmg file and authorize the application via Apple menu → System Settings → Privacy & Security → Open Anyway.

A standalone Windows executable is also available; it handles 24-hour videos but runs more slowly than the macOS build.

Alternatively, a Python source version of the Crunchometer is available on GitHub:https://github.com/RanierLabNeurobiologyAppetite2026/TheCrunchometerPy

We recommend directly downloading the standalone software from GitHub using the following link:

MacOS Silicon: **TheCrunchometer-2.0.0-macOS-arm64.dmg**

To install, double-click the .dmg file and authorize the application via Apple menu → System Settings → Privacy & Security → Open Anyway.

Windows: **Setup_TheCrunchometer-2.0.0-Windows-x64.exe**

An example folder, CrunchometerRunExample, is also provided, containing Audio_Data (.ogg) and Video_Data (.mkv) subfolders from a 10-minute recording for testing the Crunchometer software.

A tutorial video on installing and running the Crunchometer is available at the following link.

## Supplementary information

**Video 1.** Preparation of food pellets

**Video 2.** Change in HFD preference during acute semaglutide administration.

**Video 3.** Chemogenetic activation of LH GABAergic neurons induced a stress-like feeding pattern.

**Video 4.** Food Hoarding event during LH multichannel recordings.

**Supplementary Fig. 1-1.** Bill of materials

**Supplementary Fig. 1-2.** Robustness of Crunchometer bite detection to additive white noise.

**Supplementary Fig. 5-1.** Chemogenetic activation of GABAergic neurons in LH promotes eating consummatory behaviors in Unilateral and Bilateral DREADD activation.

**Supplementary Fig. 6-1**. Schematic diagram for 1 Hz pulse generator to blink an LED in synchronization with the video.

