## Supplementary Fig. 1-1 for "The Crunchometer: A Low-Cost, Open-Source Acoustic Analysis of Feeding Microstructure"

**Supplementary Fig. 1-1. Bill of materials**

| Crunchometer Bill of Materials | | | | | | | |
|---|---|---|---|---|---|---|---|
| **Main components of Crunchometer** | | | | | | | |
| **Ítem** | **Description** | **Image** | **Quantity required** | **Unit price USD** | **Webpage seller** | **Model** | **Seller** |
| **1** | **Condenser Microphone with USB Cable** | 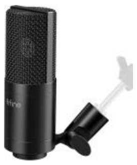 | **1** | **$ 63.88** | **https://a.co/d/evaasBZ (Accessed 03-07-2025)** | **K669** | **FIFINE** |
| **2** | **Global Shutter High Speed 120fps at 1280 x 720p USB Camera with Mini Case with CS mount 2.8-12mm varifocal lens** | 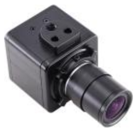 | **1** | **$ 107.24** | **https://a.co/d/22F4TgV (Accessed 03-07-2025)** | **KYT-U100-MCS2812GS1** | **Kayeton** |
| **3** | **Kimble Phenolic Black Screw Cap with Solid PE Liners, Cap Size 22-400 (Case of 144)** | 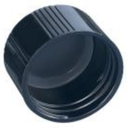 | **Bottle cap number depends on the number of pellets measured by the Crunchometer** | **$39.76** | **https://www.amazon.com/-/es/Kimble-fen%C3%B3lico-s%C3%B3lidos-polietileno-paquete/dp/B009EGMU8G?th=1** | **N/A** | **N/A** |

| | | | | | | | |
|---|---|---|---|---|---|---|---|
| | | | | | (Accessed 13-11-2025) | | |
| 4 | UHU Patafix Glue Pad Removable Reusable White | 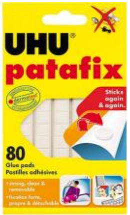 | 1 | ~$8-10 | https://www.amazon.com/dp/B01DNWS4YK<br><br>(Accessed 25-07-2025) | N/A | N/A |
| 5 | Acoustic Panels 12x12x2 Inches Black Pyramid Acoustic Foam | 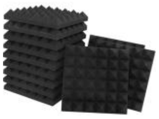 | 1 package with 12 packs | ~$19-20 | https://www.amazon.com/dp/B0928PWKH6/ref=sspa_dk_detail_right_aax_0?sp_csd=d2lkZ2V0TmFtZT1zcF9kZXRhaWxfdGhlbWF0aWNfc2hhcmVk&th=1<br><br>(Accessed 18-10-2025) | N/A | N/A |
| 6 | Acrylic Box | 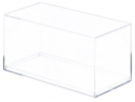 | 1 | N/A | N/A | N/A | N/A |
| **Optional: record liquid consumption (MED-PC required)** | | | | | | | |

| | | | | | | | |
|---|---|---|---|---|---|---|---|
| **7** | **Contact lickometer controller with a 28 VDC controlled MED output** | 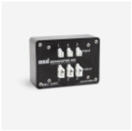 | **1** | **~$350 - 400** | **- lickometer-controller (Accessed 3-07-2025)** | **ENV-250C** | **Med Associates inc** |
| **8** | **ATmega2560 microcontroller board with USB cable.** | 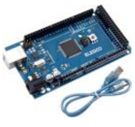 | **1** | **~$ 29.4** | **https://a.cod/d/B01H4ZLZLQ (Accessed 3-07-2025)** | **Mega 2560 R3** | **ELEGOO** |
