## Supplementary Fig. 1-2 for "The Crunchometer: A Low-Cost, Open-Source Acoustic Analysis of Feeding Microstructure"

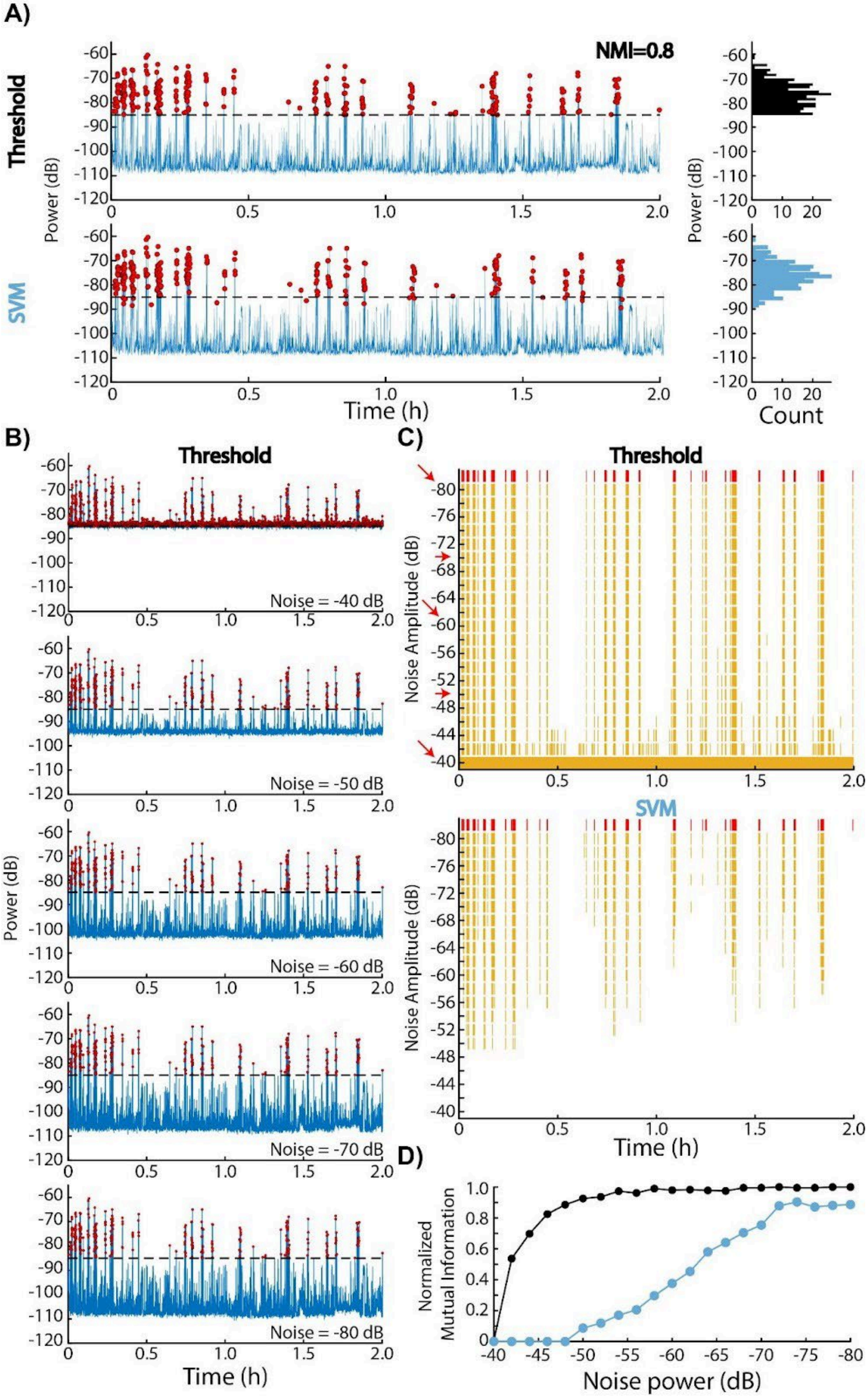

A)
Threshold
SVM
NMI=0.8
Power (dB)
Time (h)
Count
B)
Threshold
Noise = -40 dB
Noise = -50 dB
Noise = -60 dB
Noise = -70 dB
Noise = -80 dB
C)
Threshold
SVM
Noise Amplitude (dB)
D)
Normalized Mutual Information
Noise power (dB)

**Supplementary Fig 1-2. Robustness of Crunchometer bite detection to additive white noise.** The threshold-based method is sensitive to loud sounds overlapping the bite-related frequency band (500–950 Hz) but remains more reliable than the SVM-based method across the tested signal-to-noise ratio (SNR) range. **A.** Feeding detection (red dots) in a representative recording session (blue line) using the Threshold and SVM methods. Although the threshold method relies on signal amplitude within a specific frequency band (500-900 Hz), and the SVM method uses spectral pattern classification, both approaches exhibit high correlation in detecting feeding behavior, with normalized mutual information (NMI) = 0.8. **B.** Representative examples from a single recording session with increasing levels of white noise added to the original audio signal. As noise amplitude decreases, the number of detections produced by the threshold method is reduced. **C.** Ethograms of Threshold and SVM detections (top and bottom panels, respectively) across different white-noise amplitudes. Red marks indicate detections in the original audio signal, whereas yellow marks indicate detections after noise addition. Red arrows correspond to the noise levels in panel **B**. Normalized mutual information (NMI) as a function of white-noise amplitude for the Threshold and SVM methods (black and blue lines, respectively). Values near 1 indicate that the signals are identical. Increasing noise amplitude leads to more detections by the Threshold method, indicating a higher false-positive rate. In contrast, SVM detections decrease as noise amplitude increases, indicating fewer true positives. Overall, the Threshold method is more robust to white noise when detecting feeding behavior than the SVM method.
