## Supplementary Fig. 5-1 for "The Crunchometer: A Low-Cost, Open-Source Acoustic Analysis of Feeding Microstructure"

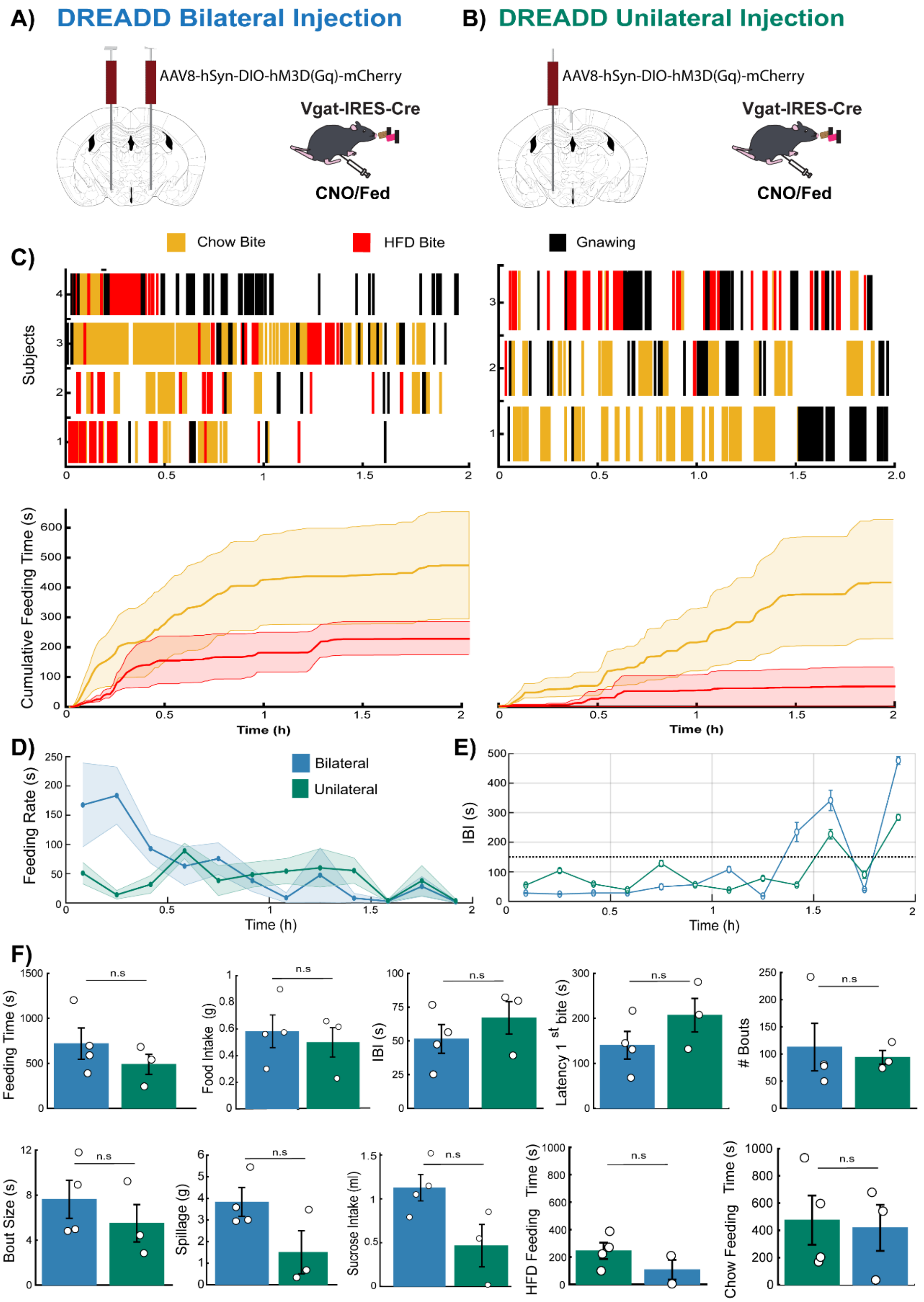


**Supplementary Fig 5-1. Chemogenetic activation of GABAergic neurons in LH promotes eating consummatory behaviors in Unilateral and Bilateral DREADD activation.** Schematic of hM3D(Gq) viral infection and experimental design. **A**) Bilateral DREADD injections into the LH of both hemispheres of fed mice (n=4) and **B**) unilateral DREADD injections into a single hemisphere (n=3). All mice were fed ad libitum and treated with clozapine-N-oxide (CNO) via intraperitoneal injection. **C)** Feeding ethograms illustrating feeding behavior following CNO administration. Red lines indicate high-fat diet

(HFD) pellet consumption, yellow lines indicate chow pellet consumption, and black lines denote gnawing events. Cumulative feeding time plots display group means (solid lines), with shaded areas representing ± SEM. **D)** Average feeding rate and **E)** inter-bite intervals (IBIs), calculated in 10-min bins across the session (top and middle panels, respectively). **(F)** Quantitative analysis of feeding behavior revealed no significant differences between unilateral and bilateral activation, including total feeding time ($t_{(5)}$ = 0.9946, $p$ = 0.3656), total food intake ($t_{(5)}$ = 0.4384, $p$ = 0.6794), spillage ($t_{(5)}$ = 2.2268, $p$ = 0.0765), number of feeding bouts ($t_{(5)}$ = 0.3536, p = 0.7381), bout size ($t_{(5)}$ = 0.8306, $p$ = 0.4440), latency to first bite ($t_{(5)}$ = 1.3080, p = 0.2478), IBIs ($t_{(5)}$ = 0.9085, p = 0.4053), intake of a 10% sucrose solution ($t_{(5)}$ = 2.4336, p = 0.0591), total chow feeding time ($t_{(5)}$ = 0.2111, p = 0.8411), and total HFD feeding time ($t_{(5)}$ = 1.9036, p = 0.1153). Data are presented as mean ± SEM. *p < 0.05; n.s., p > 0.05.
