## Supplementary Fig. 6-1 for "The Crunchometer: A Low-Cost, Open-Source Acoustic Analysis of Feeding Microstructure"

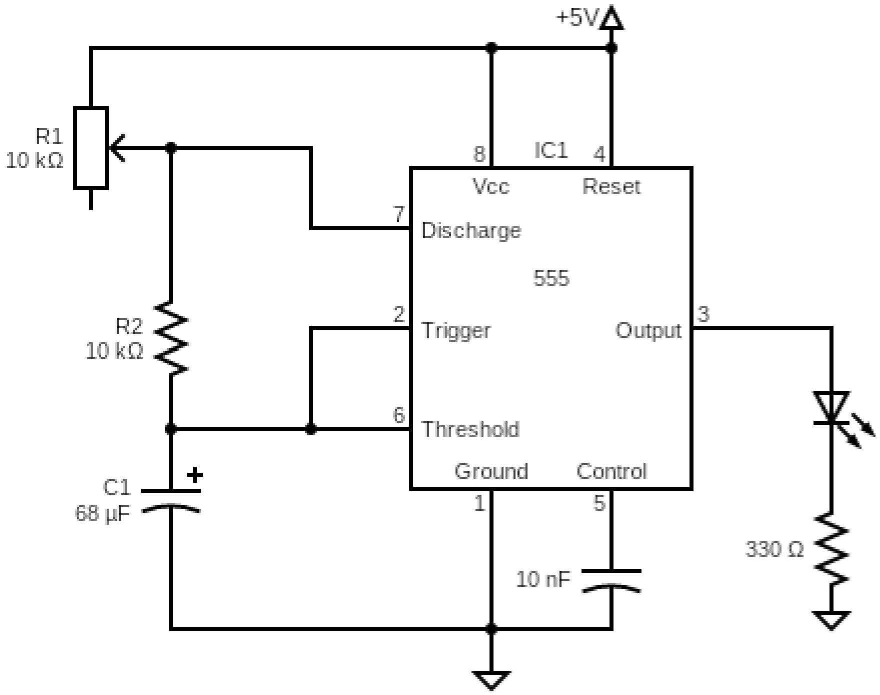


**Supplementary Fig 6-1. Schematic diagram for 1 Hz pulse generator to blink an LED in synchronization with the video.** The schematic diagram for the pulse generator uses a 555 timer in astable mode, where the resistance values of R1 and R2 and the capacitance of C1 determine the continuous square-wave frequency by considering the charge and discharge times of the capacitor (C1) through R1 and R2. To calculate the frequency of the pulse-generator the following formula is used:

frequency=1.44/0.693*R1+2R2*C1

Thus, to generate a 1Hz square wave, we used a 68 $\mu$F capacitor (C1), a 10 K$\Omega$ resistor (R2), and 10 K$\Omega$ potentiometer (R1) to adjust the frequency due to the variability in component values (e.g., tolerance).
